# Extracellular CIRP induces neurotoxic astrocytes via TREM-1 in Alzheimer’s disease

**DOI:** 10.64898/2026.09.18.752724

**Authors:** Archna Sharma, Dilara Aylar, Dmitriy Lapin, Philippe Marambaud, Ping Wang

## Abstract

Extracellular cold-inducible RNA-binding protein (eCIRP) is a crucial neuroinflammatory mediator in ischemic stroke and alcohol-induced memory impairment. We have recently discovered that amyloid-β causes microglia to release eCIRP, prompting us to investigate its role in Alzheimer’s disease (AD). We found that eCIRP levels were significantly elevated in the cerebrospinal fluid (CSF) and plasma of AD patients compared with age-matched non-AD subjects, as well as in hTau.P301S mice (a model of AD tauopathy) compared with wildtype control mice. Plasma eCIRP strongly correlated with astrocyte activation marker glial fibrillary acidic protein (GFAP) in AD patients. eCIRP induced neurotoxic astrocyte-specific genes and astrocyte release of proinflammatory and neurotoxic factors in C8-D1a cells, primary murine astrocytes and intracerebroventricular eCIRP-injected C57BL/6 mice brains. In particular, eCIRP increased Complement 3, an astrocytic marker involved in neuroinflammation-associated neurodegeneration. Primary astrocytes from TREM-1 knockout mice were resistant to eCIRP induction of neurotoxic astrocytes. eCIRP increased TREM-1 expression and activation in astrocytes. Blocking CIRP/TREM-1 interaction using peptide M3, effectively attenuated eCIRP’s induction of neurotoxic astrocytes. Thus, eCIRP strongly correlates with astrocyte reactivity in AD patients, and eCIRP induces neurotoxic astrocytes via TREM-1, which is attenuated by M3, suggesting a novel therapeutic opportunity to target neurotoxic astrocytes in AD.

## Introduction

Alzheimer’s disease (AD) is the most common form of neurodegenerative dementia and the fifth leading cause of death among people aged 65 and older, afflicting 7.4 million people in the US alone (1). Neuropathologically, AD is characterized by plaques containing aggregated amyloid β (Aβ) and neurofibrillary tangles containing abnormally phosphorylated tau (2). Tau pathology, in particular, is strongly correlated with neurodegeneration and cognitive deficits in AD (3, 4). AD pathogenesis includes neuroinflammation due to activation of microglia and astrocytes by pathological triggers such as protein aggregates (5, 6).

Astrocytes are the most abundant cell type in the CNS and have ability to initiate and amplify immune response (7). However, astrocytes are heterogenous in their reactivity and have diverse molecular profiles (8–10). Reactive astrocytes appear to correlate with AD neurodegeneration and targeting astrogliosis was proposed to be beneficial in AD cognitive deficits (11–16). Subsets of reactive astrocytes are proinflammatory and neurotoxic (17–19). While neurotoxic astrocytes are reported in AD (17, 20), its pathogenic role in AD remains elusive. There is only indirect evidence suggesting the participation of neurotoxic astrocytes in AD pathogenesis (18). Thus, demonstrating how neurotoxic astrocytes are induced in AD, and the pathways involved are critically important to developing novel strategies to prevent and treat AD.

Cold-inducible RNA-binding protein (CIRP) is a constitutively expressed nuclear regulator of protein translation which is translocated to the cytoplasm and released to the extracellular space under stress (21). Extracellular CIRP (eCIRP) acts as a danger-associated molecular pattern (DAMP) to increase tissue injury and mortality after inflammation (21–27). eCIRP is typically released by damaged neurons and activated microglial cells in response to cerebral ischemia and alcohol exposure and is a critical mediator of inflammation and injury during ischemic brain stroke (28, 29) and alcohol-induced memory impairment (24, 30, 31). Additionally, eCIRP is released by microglial cells in response to AD-associated neuronal Aβ (32). eCIRP is proinflammatory and promotes type 1 inflammation in macrophages (33, 34), neutrophils (35, 36), T lymphocytes (37–39), and lung alveolar epithelial cells (40). Triggering receptor expressed on myeloid cells-1 (TREM-1) has been implicated in the pathology of various CNS diseases including AD (41–44) and is known to induce neuroinflammation (33–37). eCIRP activates TREM-1 proinflammatory signaling on macrophages (45, 46) and other cells (40, 47–49). However, no studies have yet investigated whether eCIRP is released in AD, its effects on astrocyte reactivity, or the mechanisms involved.

Therefore, we hypothesized that eCIRP plays a major role in inducing neurotoxic astrocytes in AD. In this study, we measured eCIRP levels in the cerebrospinal fluid (CSF) and plasma of AD patients as well as in hTau.P301S mice (a model of AD tauopathy) and assessed its association with glial fibrillary acidic protein (GFAP), a marker of astrocyte activation (38). We addressed if eCIRP can induce neurotoxic astrocytes releasing proinflammatory and neurotoxic factors. As TREM-1 is an eCIRP receptor, we investigated if neurotoxic astrocyte induction could be a result of eCIRP’s direct binding to TREM-1. Finally, we tested if M3, a small anti-eCIRP inhibitory peptide blocking its interaction with TREM-1, can abolish eCIRP’s induction of neurotoxic astrocytes. Herein we have provided strong evidence in support of elevated eCIRP levels in AD patients and eCIRP’s induction of neurotoxic astrocytes. We have identified the receptor and signaling involved and potential inhibitor attenuating eCIRP-induced neurotoxic astrocytes.

## Results

### eCIRP levels are elevated in AD patients and AD-tauopathy mice and correlate with GFAP

Since AD is associated with blood-brain barrier disruption leading to increased plasma levels of Aβ, p-tau181, and GFAP (50), we evaluated whether levels of eCIRP are elevated in the blood of AD patients compared to age- and sex-matched non-AD control subjects (thirty-seven subjects per group with 14 males and 23 females in each group). The eCIRP levels in the plasma of AD patients were 5.1-fold higher than those in control subjects (**Figure 1A**). Of note, the plasma levels of eCIRP in female AD patients were 1.4-fold higher than that of male AD patients (**Supplementary Figure 1**). We also measured eCIRP in the cerebrospinal fluid (CSF) of twelve AD patients and twelve age-matched unaffected control subjects obtained from the Feinstein Institutes’ Litwin-Zucker Center for Alzheimer’s Disease Research repository and diagnosed according to current criteria (51, 52). The CSF eCIRP levels in AD patients were 1.5-fold higher than those in control subjects (**Figure 1B**). This clinical finding supports our hypothesis that eCIRP plays an important role in AD pathogenesis.

**Figure 1.**
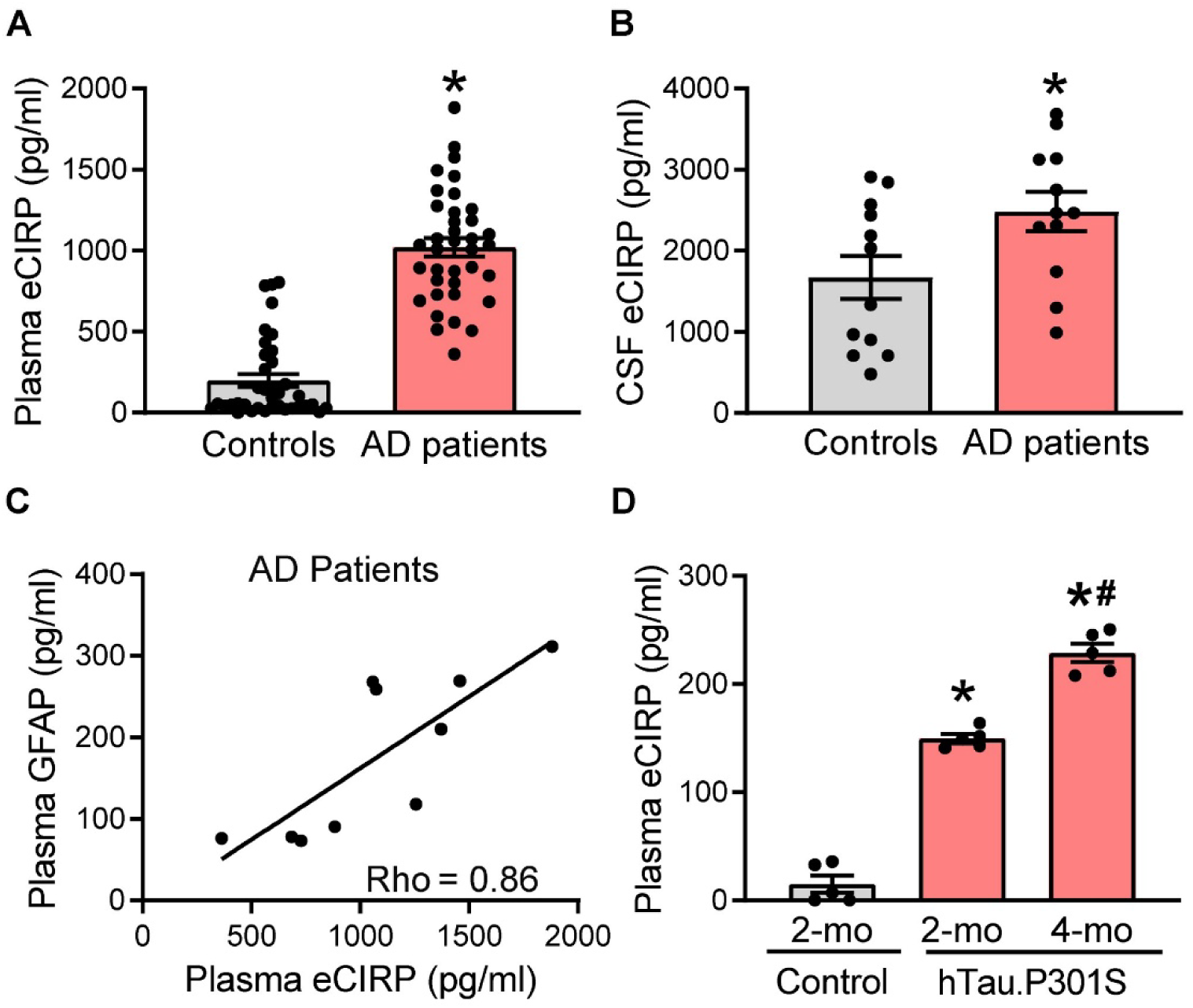
eCIRP levels are elevated in AD patients as well as AD-tauopathy mice and plasma eCIRP and GFAP correlate in AD patients. (**A**) The plasma levels of eCIRP were measured by ELISA in AD patient and age- and sex-matched unaffected control samples obtained from Precision Biospecimen Solutions (Bethesda, MD). There are 14 males (37.84%) and 23 females (62.16%) per group with the age range 63-88 years and no significant difference in median age (77 years for control group and 79 years for AD group). Experiments were performed at least 3 times, and all data were analyzed. Data are expressed as means ± SEM and were compared using Mann-Whitney test. n = 37 per group; *p < 0.0001. (**B**) The CSF levels of eCIRP were measured by ELISA in de-identified AD patients and unaffected control samples obtained from the Litwin-Zucker Center for Alzheimer’s Disease Research repository. Data are expressed as means ± SEM and were compared using Mann-Whitney test. n = 12 per group; *p < 0.05. (**C**) The plasma levels of eCIRP and astrocyte activation marker GFAP were measured by ELISA in AD patient samples obtained from Precision Biospecimen Solutions (Bethesda, MD). There are 5 AD males (50%) and 5 AD females (50%) with the age range 63-88 years and median age of 72 years. Data are expressed as means ± SEM and correlation was analyzed using Linear Regression and Spearman’s Correlation. n = 10 per group; R^2^ = 0.63 and \**p* = 0.006 for Linear Regression Analysis, Rho = 0.86 and \**p* = 0.003 for Spearman’s Correlation. (**D**) The plasma levels of eCIRP in homozygous 2-month-old and 4-month-old male hTau.P301S mice and 2-month-old male C57BL/6 (control) mice were measured by ELISA. Data are expressed as means ± SEM and were analyzed using one-way ANOVA, Tukey’s multiple comparisons test. n = 5 per group; *p < 0.0001 vs. control, ^#^p < 0.0001 vs. 2-month-old hTau.P301S mice.

The astrogliosis marker glial fibrillary acidic protein (GFAP) is elevated in the blood in AD patients and associated with AD cognitive impairment (53, 54). In AD, GFAP increases in plasma earlier than in the CSF (55). Therefore, we measured levels of GFAP and eCIRP in ten plasma samples from AD patients (five of each sex). eCIRP and GFAP levels significantly showed strong positive correlation in the AD patient’s plasma cohort tested (**Figure 1C**). This correlation finding strongly suggests that eCIRP may promote astrocyte reactivity in AD patients. Next, we determined eCIRP levels in hTau.P301S mice, which overexpress human tau mutant P301S under the control of murine neuron-specific promoter Thy-1 (56) and develop neuropathology and behavioral deficits by 4 months of age (57). eCIRP levels in the blood of 2-month-old homozygous hTau.P301S mice (i.e., prior to the appearance of tau tangles) were 9.9-fold higher than that of control mice (**Figure 1D**). Moreover, plasma eCIRP levels further increased with progression in tau pathology with 1.5-fold increase at 4-months (i.e., after disease onset) compared to 2-month-old hTau.P301S mice (**Figure 1D**). The increased levels of eCIRP in AD patients as well as in hTau.P301S mice prior to disease onset suggest that eCIRP may play a pathogenic role in the development of AD. Taken together, these findings point to eCIRP’s potential role in AD pathogenesis via regulating astrocyte reactivity.

### eCIRP induces neurotoxic astrocytes

We first verified that eCIRP significantly increased the mRNA expression of reactive astrocyte activation marker GFAP in C8-D1a cells as well as magnetically purified primary astrocytes (**Supplementa**ry **Figure 2A**). Of note, eCIRP also induced mRNA expression of ALDH1, which is a pan-astrocytic marker of astrocytic differentiation, in C8-D1a cells as well as primary astrocytes (**Supplementa**ry **Figure 2B**). GFAP protein expression and eCIRP-induced GFAP increase were also confirmed in primary astrocytes (**Supplementary Figure 3**). To determine whether eCIRP can induce neurotoxic astrocyte differentiation, we stimulated the mouse astrocytic cell line C8-D1a with eCIRP at varying doses (0-2.5 µg/ml) for variable times (0-24 hours). We assessed the mRNA expression of neurotoxicity-associated astrocyte marker Complement 3 (C3), neuroprotection-associated astrocyte marker S100 calcium binding protein A10 (S100A10), and inflammatory cytokine interleukin-6 (IL-6) release in C8-D1a cells. eCIRP significantly increased the mRNA expression of C3 in C8-D1a cells in time- and dose-dependent manner with a significant 3-fold increase at 24 hours (**Figure 2A**) and 2.1–fold increase with 2.5 µg/ml eCIRP (**Figure 2B**). Interestingly, eCIRP did not alter the mRNA expression of the neuroprotection-associated marker S100A10 (**Figure 2A-B**). In addition, eCIRP time- and dose-dependently increased the IL-6 release from C8-D1a cells with a significant 1.71-fold increase at 24 hours (**Figure 2C**) and 1.65-fold increase with 2.5 µg/ml eCIRP (**Figure 2D**). Next, we evaluated the protein expression of C3 and S100A10 as well as the release of IL-6 and neurotoxic factor lipocalin 2 (Lcn2) in C8-D1a cells and primary mouse astrocytes stimulated with 2.5 µg/ml eCIRP for 24 hours. As expected, eCIRP also increased C3 protein levels by 2.2-fold in C8-D1a cells and by 16.8-fold in primary astrocytes (**Figure 2E**), whereas there were no significant changes in S100A10 protein levels in either C8-D1a cells or primary astrocytes (**Figure 2F**). eCIRP also increased IL-6 release in the conditioned medium by 1.9-fold for C8-D1a cells and by 35.8-fold for primary astrocytes (**Figure 2G**). Furthermore, eCIRP increased Lcn2 release by 9.9-fold in C8-D1a cells conditioned medium. (**Figure 2H**). While unstimulated control primary astrocytes did not release any Lcn2, with eCIRP stimulation primary astrocytes induced release of 1629 ± 104 pg/ml Lcn2 (**Figure 2H**). Primary astrocytes consistently showed higher response to eCIRP stimulation compared to C8-D1a cells.

**Figure 2.**
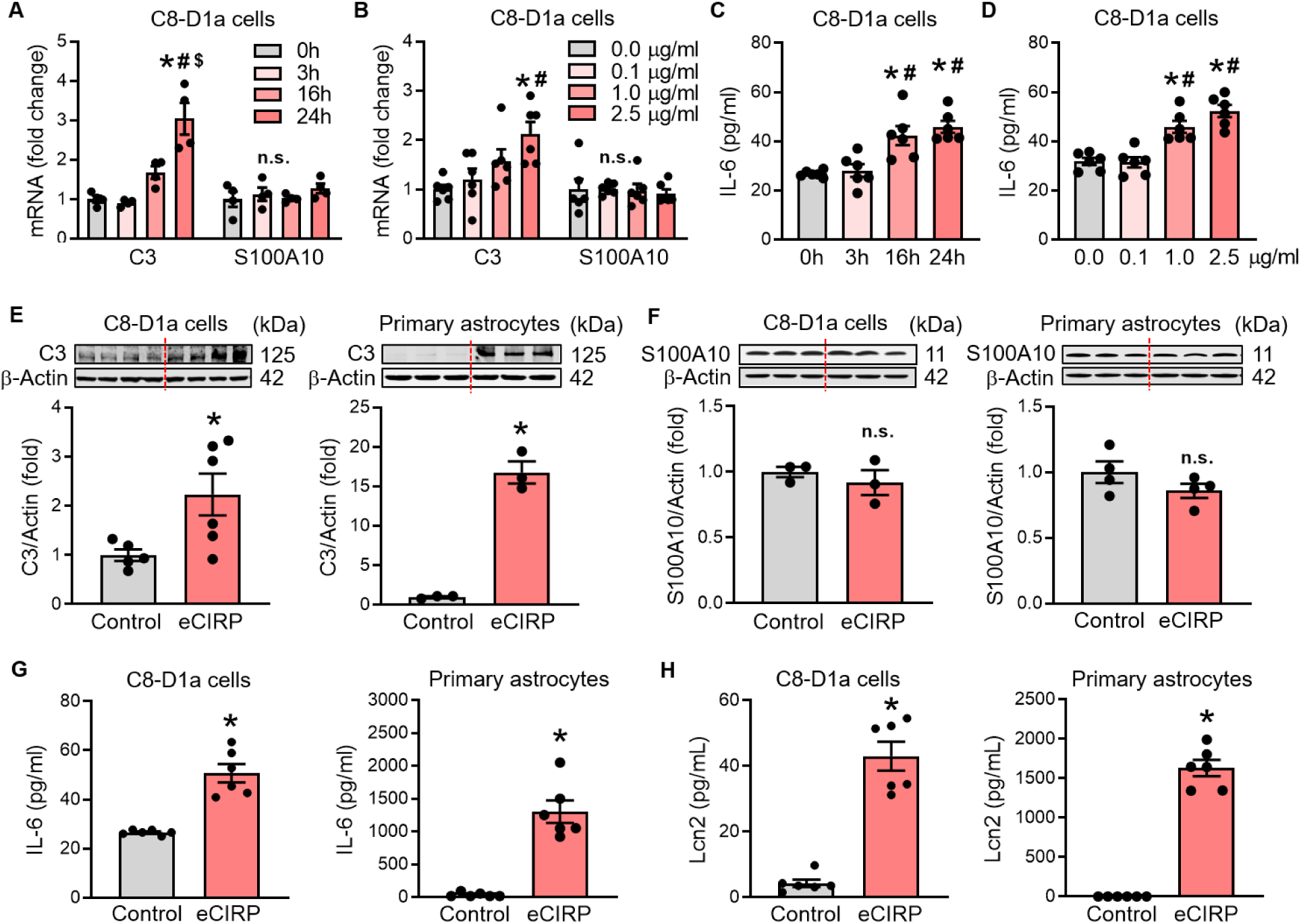
eCIRP induces neurotoxic astrocytes in time- and dose-dependent manner. (**A-D**) C8-D1a cells (0.3 × 10^6^/ml) were treated with recombinant mouse CIRP (eCIRP) at indicated times or doses. (**A**) RT-qPCR quantification of relative C3 and S100A10 gene expression in C8-D1a cells treated with 1 µg/ml eCIRP at indicated times. Data are expressed as means ± SEM and were compared using two-way ANOVA, Sidak’s multiple comparisons test. n = 4 per group; *p < 0.0001 vs. 0 hour, ^#^ p < 0.0001 vs. 3 hours, ^$^ p = 0.0003 vs. 16 hours, n.s. = 0.9. (**B**) RT-qPCR quantification of relative C3 and S100A10 gene expression in C8-D1a cells treated with eCIRP for 24 hours at indicated doses. Data are expressed as means ± SEM and were compared using two-way ANOVA, Sidak’s multiple comparisons test. n = 6 per group; *p = 0.0003 vs. no eCIRP (0 µg/ml), ^#^ p = 0.0027 vs. 0.1 µg/ml eCIRP, n.s. = 0.9. (**C**) IL-6 released in the conditioned medium of C8-D1a cells treated with 1 µg/ml eCIRP at indicated times quantified by ELISA. Data are expressed as means ± SEM and were compared using one-way ANOVA, Tukey’s multiple comparisons test. n = 6 per group; *p < 0.003 vs. 0 hour, ^#^ p < 0.007 vs. 3 hours. (**D**) IL-6 released in the conditioned medium of C8-D1a cells treated with eCIRP for 24 hours at indicated doses quantified by ELISA. Data are expressed as means ± SEM and were compared using one-way ANOVA, Tukey’s multiple comparisons test. n = 6 per group; *p < 0.001 vs. no eCIRP (0 µg/ml), ^#^ p < 0.001 vs. 0.1 µg/ml eCIRP. (**E-H**) C8-D1a cells (0.3 × 10^6^/ml) or primary astrocytes (0.3 × 10^6^/ml) were treated with 2.5 µg/ml eCIRP for 24 hours. Representative Western blot images and bar graph from the densitometric analysis of (**E**) C3 and (**F**) S100A10 immunoblots for C8-D1a cells and primary astrocytes as indicated. The dotted lines on the Western blot image reflect different samples from the corresponding groups shown in the bar graph below. Data are expressed as means ± SEM and were compared by Unpaired *t*-test. n = 3-6 per group; * p < 0.05 vs. control (no eCIRP), n.s. = 0.5 for C8-D1a cells and n.s. = 0.2 for primary astrocytes. IL-6 (**G**) and Lcn2 (**H**) released in the conditioned medium of C8-D1a cells and primary astrocytes as indicated quantified by ELISA. Data are expressed as means ± SEM and were compared by Unpaired *t*-test. n = 6 per group; * p < 0.0001 vs. control (no eCIRP).

To further confirm eCIRP-induction of neurotoxic gene signatures in astrocytes, we assessed the mRNA expression of a larger panel of astrocyte reactivity markers (17) in C8-D1a cells and primary astrocytes stimulated with 2.5 µg/ml eCIRP for 24 hours. eCIRP significantly increased the expression of all five neurotoxicity-associated genes (C3, H2-T23, H2-D1, GBP2 and PSMB8) in C8-D1a cells (**Figure 3A**) as well as in primary astrocytes (**Figure 3B**). To evaluate whether this also occurred *in vivo*, we intracerebroventricularly (*icv)*-injected 1 µg eCIRP into the brain of 10-week-old C57BL/6 mice and 24 hours later harvested the whole brain tissue for RNA extraction. *icv*-injected eCIRP also significantly increased brain mRNA expression of all five neurotoxicity-associated genes (**Figure 3C**). Interestingly, the *icv*-eCIRP injected mice brains *in vivo* induced the neurotoxic signature genes at very similar levels as the primary astrocytes stimulated with eCIRP *in vitro* (**Figure 3B-C**). In addition, eCIRP did not alter the expression of any of the five neuroprotection-associated genes (S100A10, CLCF1, EMP1, SLC10A6 and B3GNT5) *in vitro* in C8-D1a cells (**Figure 3A**) and primary astrocytes (**Figure 3B**) nor *in vivo* in *icv*-eCIRP injected mice brains (**Figure 3C**). These results demonstrate, for the first time, that eCIRP is able and sufficient to induce the expression of neurotoxic gene signature on astrocytes.

**Figure 3.**
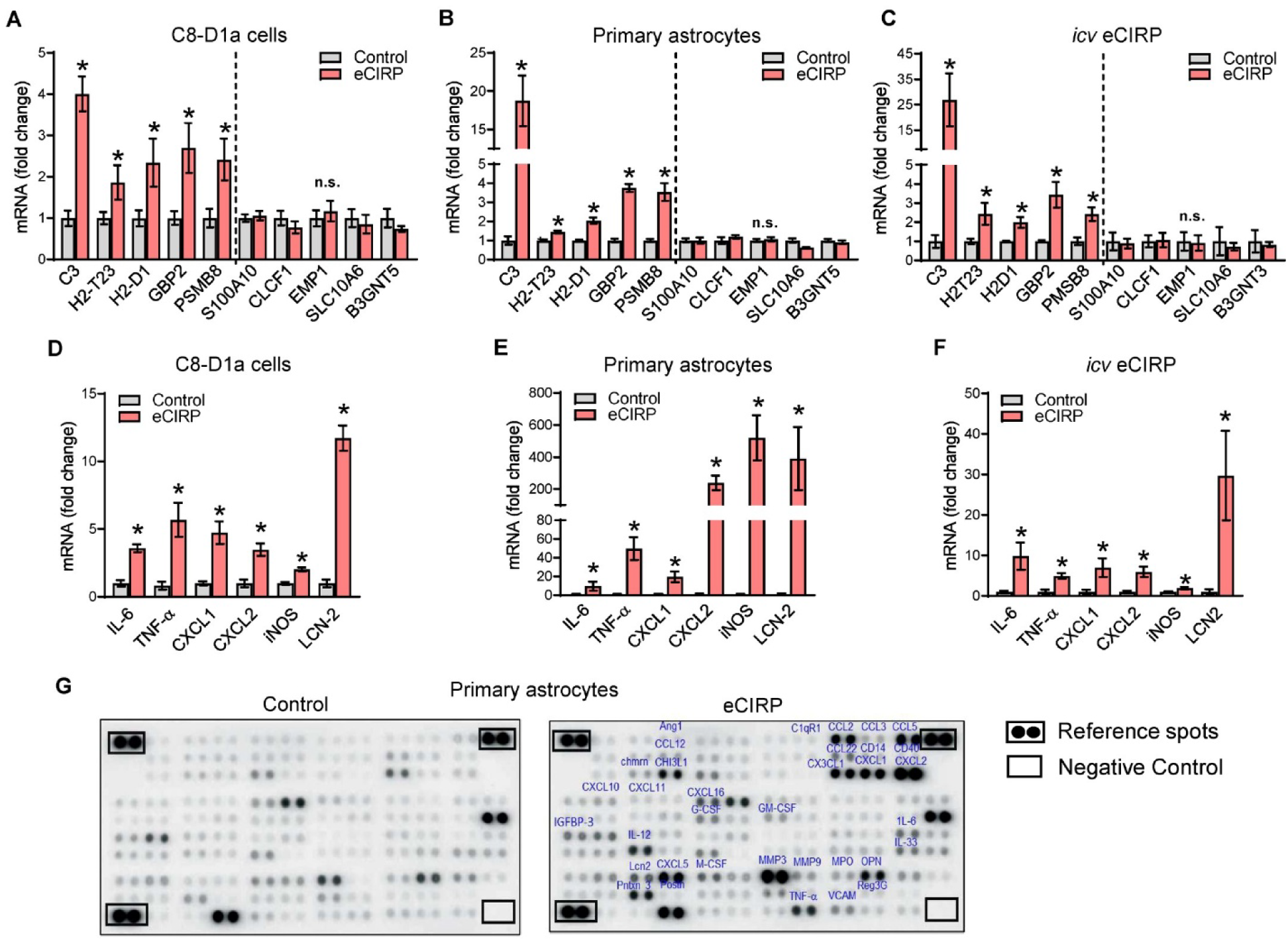
eCIRP induces neurotoxic astrocyte gene signature along with inflammatory and neurotoxic mediators in murine astrocytes. **(A-B)** C8-D1a cells (0.3 × 10^6^/ml) or primary astrocytes (0.3 × 10 /ml) were treated with 2.5 µg/ml eCIRP for 24 hours. RT-qPCR quantification of the relative mRNA expression of astrocyte-reactivity signature genes panel (including C3, H2-T23, H2-D1, GBP2, PSMB8, S100A10, CLCF1, EMP1, SLC10A6 and B3GNT5) *in vitro* in (**A**) C8-D1a cells and (**B**) primary astrocytes. Data are expressed as means ± SEM and were compared using Multiple group *t-*test. n = 10-12 per group for (**A**) and n = 5-6 per group for (**B**); * p < 0.05 vs. control (no eCIRP), n.s., not significant p > 0.05. (**C**) C57BL/6 mice were *icv*-injected with eCIRP (1 μg) or PBS and the brains harvested 24 hours later. RT-qPCR quantification of the relative mRNA expression of the same ten astrocyte reactivity signature gene panel *in vivo* in the whole brains. Data are expressed as means ± SEM and were compared using Multiple group *t-*test. n = 5 per group; * p < 0.05 vs. control (no eCIRP), n.s., not significant p > 0.05. (**D-E**) C8-D1a cells (0.3 × 10 /ml) or primary astrocytes (0.3 × 10 /ml) were treated with 2.5 µg/ml eCIRP for 24 hours. RT-qPCR quantification of the relative mRNA expression of inflammatory cytokines (TNF-α, IL-6), chemokines (CXCL1, CXCL2), inducible nitric oxide (iNOS) and neurotoxic factor Lcn2 *in vitro* in (**D**) C8-D1a cells and (**E**) primary astrocytes. Data are expressed as means ± SEM and were compared using Multiple group *t-*test. n = 6 per group for (**D**) and n = 5-6 per group for (**E**); * p < 0.05 vs. control (no eCIRP), n.s., not significant p > 0.05. (**F**) C57BL/6 mice were *icv*-injected with eCIRP (1 μg) or PBS and the brains harvested 24 hours later. RT-qPCR quantification of the relative mRNA expression of the same six astrocyte inflammatory genes *in vivo* in the whole brains. Data are expressed as means ± SEM and were compared using Multiple group *t-*test. n = 4 per group; * p < 0.05 vs. control (no eCIRP), n.s., not significant p > 0.05. (**G**) The conditioned medium of primary astrocytes treated with 2.5 µg/ml eCIRP for 24 hours was screened using mouse cytokine proteome profiler for 111 inflammatory mediators released. Representative proteome profiler membrane images from control and eCIRP treated groups as indicated. Each mediator is in pair in duplicate spots and the location of the pairs of positive reference spots in three corners of each array and the pair of negative control spots on each membrane is indicated for orientation. The densities of the spots on the images of both membranes were quantified and compared. Thirty-five inflammatory mediators upregulated in the eCIRP treatment group compared to the control group are labeled on the eCIRP group membrane image.

To verify whether eCIRP-induced astrocytes are indeed inflammatory, we stimulated C8-D1a cells or primary mouse astrocytes with 2.5 µg/ml eCIRP and assessed the effects on inflammatory mediators 24 hours later. eCIRP significantly induced the mRNA expression of inflammatory cytokines (IL-6, TNF-α), chemokines (CXCL1, CXCL2), inducible nitric oxide synthase (iNOS), and neurotoxic factor lipocalin 2 (Lcn2) in C8-D1a cells (**Figure 3D**), as well as in primary astrocytes (**Figure 3E**). We also *icv*-injected eCIRP *in vivo* and, 24 hours later, eCIRP significantly increased the mRNA expression of the same inflammatory and neurotoxic mediators in the brain tissue (**Figure 3F**). Next, to further confirm the changes in mRNA expression, we assessed the level of 111 soluble inflammatory proteins in the conditioned medium of primary astrocytes. After 24 hours of stimulation, eCIRP induced astrocyte release of 35 proteins, including a variety of chemokines (CXCL1, CXCL2, CXCL5, CXCL16, CCL2, and CCL5), cytokines (IL-6, TNF-a, IL-12, and IL-33), neurotoxic factors (Lcn2 and IGFBP-3) and other neuroinflammatory mediators (MMP3, CHI3L1, M-CSF, G-CSF and GM-CSF, and OPN) (**Figure 3G**). Since eCIRP induces proinflammatory and neurotoxic astrocytes, the high levels of eCIRP in AD patients further support that eCIRP-induced astrocyte-associated inflammation and neurotoxicity play a causal role in AD.

### TREM-1 mediates the induction of neurotoxic astrocytes by eCIRP

eCIRP has been shown to bind to TREM-1 with very high affinity and activate the downstream inflammatory signaling in macrophages (45). To confirm the role of TREM-1 in the induction of neurotoxic astrocytes by eCIRP, we stimulated magnetically purified primary astrocytes from WT and TREM-1^−/−^ mice with eCIRP for 24 hours and assessed protein expression of C3. Total cellular C3 protein levels increased with eCIRP stimulation by 11.7-fold in WT primary astrocytes, whereas that increased by only 3.1-fold in TREM-1^−/−^ astrocytes (**Figure 4A**). C3 levels in TREM-1^−/−^ astrocytes were significantly lower by 73.9% compared to WT astrocytes (**Figure 4A**). Since reactive astrocytes promote neurotoxicity via the release of C3 which causes neuronal damage via C3a receptor on neurons and microglia, we additionally assessed levels of C3 released from astrocytes. In the absence of eCIRP stimulation, WT and TREM-1^−/−^ control astrocytes did not release any C3, while TREM-1^−/−^ astrocytes showed significant downregulation by 54.1% in eCIRP-induced released C3 levels compared to WT astrocytes (**Figure 4B**). eCIRP stimulation of WT and TREM-1^−/−^ astrocytes induced upregulation of mRNA expression of neurotoxic markers C3, H2-T23 and PSMB8 (**Figure 4C**), inflammatory mediators (**Figure 4D**), as well as release of TNF-α and IL-6 (**Figure 4E**). However, eCIRP-induced mRNA expression levels in TREM-1^−/−^ astrocytes were significantly downregulated by 85.3% for C3, 77.1% for H2-T23, and 58.0% for PSMB8 compared to WT astrocytes (**Figure 4C**). TREM-1^−/−^ astrocytes also significantly downregulated mRNA expression of IL-6 by 59.0%, TNF-α by 62.8%, and iNOS by 78.6% compared to eCIRP-induced WT astrocytes (**Figure 4D**). Moreover, eCIRP-induced cytokine levels in the conditioned medium were also significantly decreased (TNF-α by 41.5% and IL-6 by 28.1%) in TREM-1^−/−^ astrocytes compared to WT astrocytes (**Figure 4E**).

**Figure 4.**
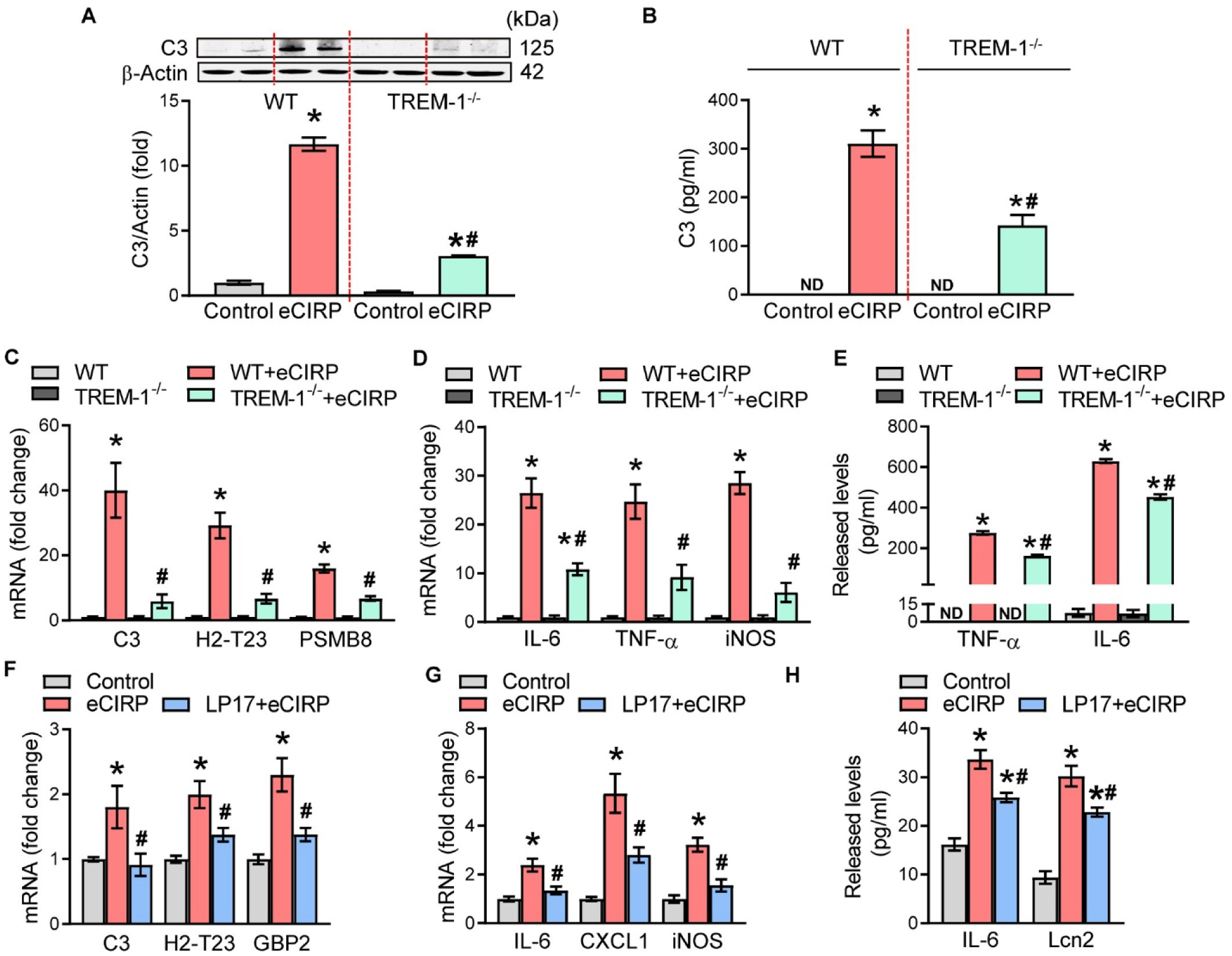
eCIRP induces neurotoxic astrocytes via TREM-1. WT or TREM-1^−/−^ primary astrocytes (0.3 × 10^6^ /ml) were treated with 2.5 µg/ml eCIRP for 24 hours. Conditioned media were harvested and cells were lysed for protein or mRNA extraction. (**A**) Representative Western blot images and bar graph from the densitometric analysis of intracellular C3 protein. The dotted lines on the Western blot image reflect different samples from the corresponding groups shown in the bar graph below. Data are expressed as means ± SEM and were compared using one-way ANOVA, Tukey’s multiple comparisons test. n = 3 per group; *p < 0.0001 vs. no eCIRP control for WT or TREM-1^−/−^,^#^p < 0.0001 vs. WT eCIRP. (**B)** Released C3 in the conditioned media was quantified by ELISA. Data are expressed as means ± SEM and were compared using one-way ANOVA, Tukey’s multiple comparisons test. n = 8-10 per group; *p < 0.0001 vs. no eCIRP control for WT or TREM-1, p < 0.0001 vs. WT eCIRP. (**C**) RT-qPCR quantification of the relative mRNA expression of C3, H2-T23, and PSMB8. Data are expressed as means ± SEM and were compared using Mixed-effects model (REML) with Tukey’s multiple comparisons test. n = 5-8 per group; * p < 0.05 vs. WT or TREM-1^−/−^ control (no eCIRP), ^#^p < 0.05 vs. WT eCIRP. (**D**) RT-qPCR quantification of the relative mRNA expression of TNF-α, IL-6, and iNOS. Data are expressed as means ± SEM and were compared using Mixed-effects model (REML) with Tukey’s multiple comparisons test. n = 5-8 per group; WT or TREM-1^−/−^ control (no eCIRP), ^#^p < 0.05 vs. WT eCIRP. (**E)** Released TNF-α and IL-6 in the conditioned media was quantified by ELISA. Data are expressed as means ± SEM and were compared using two-way ANOVA, Tukey’s multiple comparisons test. n = 6 per group; *p < 0.0001 vs. no eCIRP control for WT or TREM-^1−/−^, ^#^p < 0.0001 vs. WT eCIRP. (**F-H)** C8-D1a cells (0.3 × 10^6^ /ml) were pretreated with 80 μg/ml LP17 (TREM-1 inhibitor peptide) for 30 min followed by stimulation with 2.5 µg/ml eCIRP for 24 hours. Conditioned media were harvested and cells were lysed for mRNA extraction. RT-qPCR quantification of the relative mRNA expression of (**F)** neurotoxic astrocyte markers (C3, H2-T23, GBP2) and (**G)** inflammatory mediators (TNF-α, IL-6, iNOS). (**H**) IL-6 and Lcn2 released in the conditioned medium quantified by ELISA. Data are expressed as means ± SEM and were compared using Mixed-effects model (REML) with Tukey’s multiple comparisons test. n = 6-8 per group for (**F-G**) and n = 3-4 per group for (**H**); *p < 0.05 vs. no eCIRP control, ^#^p < 0.05 vs. eCIRP alone.

Likewise, pretreatment of C8-D1a cells with the TREM-1 inhibitor LP17 (58) significantly abrogated the eCIRP-induced mRNA expression of neurotoxic astrocyte markers C3 (by 49.3%), H2-T23 (by 31.0%) and GBP2 (by 40.0%) (**Figure 4F**), as well as inflammatory mediators IL-6 (by 43.8%), CXCL-1 (by 47.4%), and iNOS (by 51.8%) (**Figure 4G**), without affecting cell viability. LP17 also significantly lowered the eCIRP-induced astrocytic release of IL-6 (by 23.3%) and Lcn2 (by 24.4%) in C8D1 cells (**Figure 4H**). LP17 attenuation of released TNF-α levels was most significant in primary astrocytes (**Supplementary Figure 4A**), while attenuation levels for IL-6 (**Supplementary Figure 4B**) and Lcn2 (**Supplementary Figure 4C**) in primary astrocytes were similar to C8-D1a cells. These studies clearly demonstrate that TREM-1 is required for induction of neurotoxic astrocytes by eCIRP.

### eCIRP upregulates astrocytic TREM-1 expression and signaling

eCIRP induces TREM-1 expression in macrophages (45, 46) and variety of other cells (40, 47–49) and TREM-1 is also expressed in astrocytes (59). To verify whether eCIRP induces TREM-1 in astrocytes, we stimulated C8-D1a cells or primary mouse astrocytes with 2.5 µg/ml eCIRP and assessed the effects on TREM-1 expression. eCIRP significantly increased the mRNA expression of TREM-1 by 2.1-fold in C8-D1a cells and by 7.1–fold in primary astrocytes (**Figure 5A**). Next, we did immunofluorescence staining for TREM-1 in primary astrocytes and verified visually using confocal microscopy that eCIRP indeed increases astrocytic TREM-1 protein expression as well (**Figure 5B**). We further analyzed the cell surface expression of astrocytic TREM-1 using flow cytometry and demonstrated that eCIRP increased the surface TREM-1 expression by 3-fold in C8-D1a cells and by 2.3–fold in primary astrocytes (**Figure 5C**). Interestingly, eCIRP time-dependently increased total TREM-1 protein levels by 1.9-fold starting as early as 20 minutes and by 2-fold at 30 minutes (**Figure 5D**). To determine if eCIRP can activate proinflammatory signaling downstream of TREM-1 in astrocytes, we analyzed the immediate downstream mediator Syk and effector NFκB. eCIRP stimulation of C8-D1a cells for 30 minutes showed 1.4-fold higher levels of activated phospho-Syk compared to control C8-D1a cells (**Figure 5E**). Finally, eCIRP elevated NFκB mRNA levels by 3.3-fold in C8-D1a cells and by 2.1-fold in primary astrocytes (**Figure 5F**). Moreover, eCIRP-induction of NFκB mRNA starts early and, in a time-dependent manner (**Supplementary Figure 5A**). Finally, we found that eCIRP also significantly increased NFκB protein levels by 2.3-fold at 30 minutes (**Supplementary Figure 5B**). These results suggest that eCIRP upregulates TREM-1 expression and activation to induce neurotoxic astrocytes.

**Figure 5.**
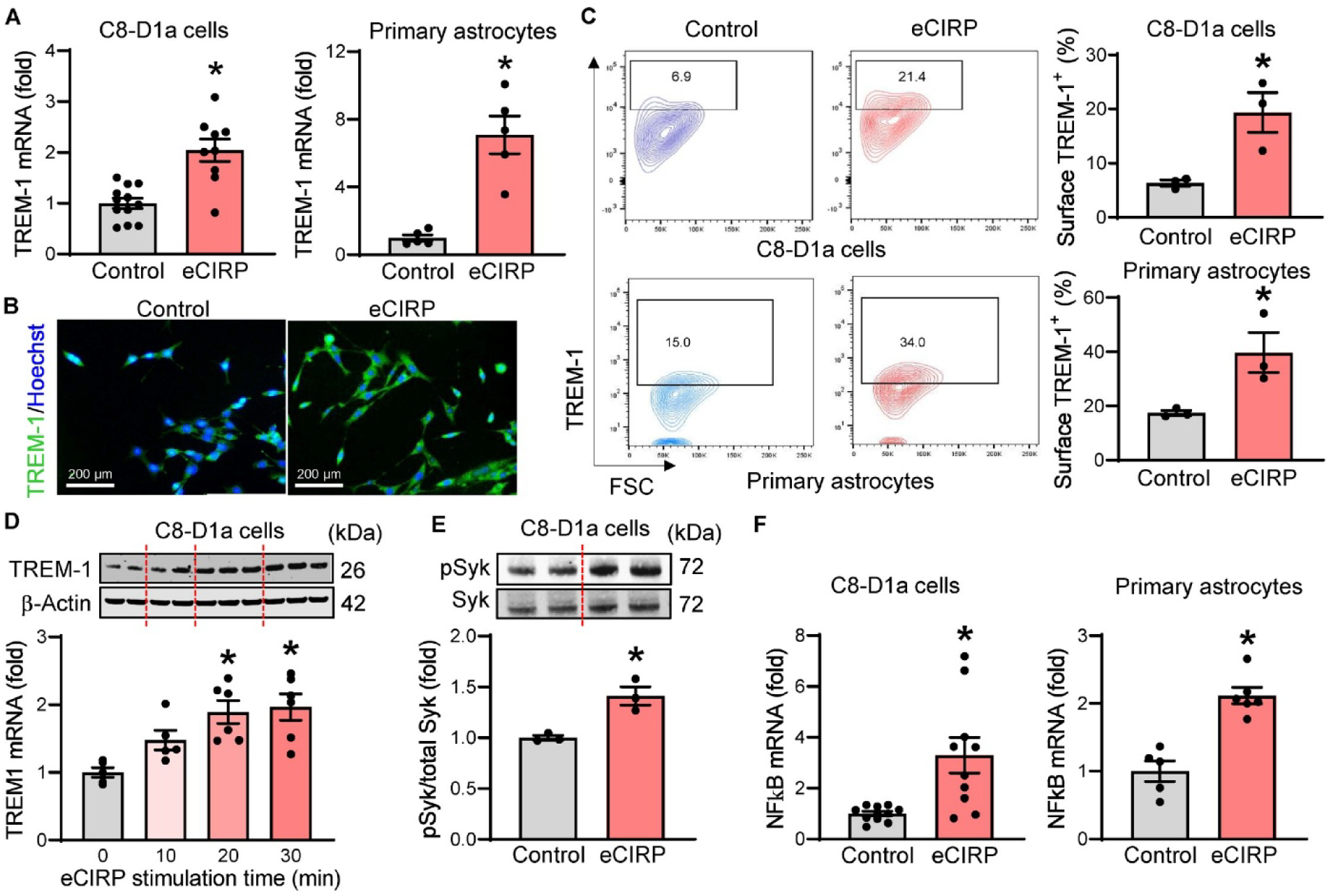
eCIRP induces TREM-1 expression and activation in murine astrocytes. **(A-C)** C8-D1a cells (0.3 × 10^6^ /ml) or primary astrocytes (0.3 × 10^6^ /ml) were treated with 2.5 µg/ml eCIRP for 24 hours and subjected to RNA extraction or staining for confocal microscopy or flow cytometry for assessing TREM-1 expression. (**A**) RT-qPCR quantification of the relative mRNA expression of TREM-1 in C8-D1a cells and primary astrocytes as indicated. Data are expressed as means ± SEM and were compared using Unpaired *t-*test. n = 10-12 per group and * p = 0.0001 vs. control (no eCIRP) for C8-D1a cells; n = 5-6 per group and * p = 0.0007 vs. control (no eCIRP) for primary astrocytes. (**B**) Representative confocal microscopy images for TREM-1 protein expression in control and eCIRP stimulated primary astrocytes stained for Hoechst (blue), and TREM-1 (green). Magnification 63x, scale bar, 200 µM. (**C**) Representative flow cytometry contour plots showing gated TREM-1^+^ astrocytes and corresponding quantification graphs showing percentages of surface TREM-1^+^ astrocytes in C8-D1a cells and primary astrocytes as indicated. Data are expressed as means ± SEM and were compared using Unpaired *t-*test. n = 3 per group for C8-D1a cells and primary astrocytes; * p < 0.05 vs. control (no eCIRP). (**D-E**) C8-D1a cells (0.3 × 10^6^ /ml) were treated with 2.5 µg/ml eCIRP for (**D**) 0, 10, 20 and 30 minutes for TREM-1 protein expression or (**E**) 30 minutes for Syk phosphorylation. Representative Western blot images and bar graph from the densitometric analysis of (**D**) TREM-1 normalized to β-actin protein and (**E**) p-Syk normalized to total Syk protein. The dotted lines on the Western blot images reflect different samples from the corresponding groups shown in the bar graph below. Data are expressed as means ± SEM and were compared using (**D**) one-way ANOVA, Tukey’s multiple comparisons test or (**E**) Unpaired *t-*test. n = 3-6 per group; *p < 0.05 vs. no eCIRP control. (**F**) C8-D1a cells (0.3 × 10^6^ /ml) or primary astrocytes (0.3 × 10^6^ /ml) were treated with 2.5 µg/ml eCIRP for 24 hours and subjected to RNA extraction. RT-qPCR quantification of the relative mRNA expression of NFκB in C8-D1a cells and primary astrocytes as indicated. Data are expressed as means ± SEM and were compared using Unpaired *t-*test. n = 10-11 per group and * p = 0.003 vs. control (no eCIRP) for C8-D1a cells; n = 5-6 per group and * p = 0.0003 vs. control (no eCIRP) for primary astrocytes.

### M3 inhibits eCIRP’s induction of neurotoxic astrocytes

To evaluate M3’s ability to inhibit eCIRP’s induction of neurotoxic astrocytes, we pretreated C8-D1a and primary astrocytes with 25 μg/ml M3 followed by stimulation with 2.5 μg/ml eCIRP for 24 hours. Without affecting cell viability, M3 abolished eCIRP induction of GFAP mRNA expression in primary astrocytes (**Supplementary Figure 6**). Indeed, M3 suppressed eCIRP-induced expression of all five neurotoxic astrocyte markers, C3 by 33.2%, H2-T23 by 29.2%, H2-D1 by 34.6%, GBP2 by 46% and PSMB8 by 35.9%, bringing most of them down to baseline control levels in C8-D1a astrocytic cells (**Figure 6A**). Similarly, M3 also abrogated eCIRP-induced mRNA expression of same neurotoxic marker genes in primary astrocytes (C3 by 67.6%, H2-T23 by 53.4%, H2-D1 by 46.6%, GBP2 by 55.8% and PSMB8 by 57.3%) (**Figure 6B**). To evaluate if M3 is also effective *in vivo*, we *icv* co-injected 10 μg/ml M3 or vehicle control with 1 µg eCIRP into the brain of 10-week-old C57BL/6 mice. At 24 hours after *icv*-injection, we harvested the whole brain tissue for RNA extraction and analyzed the expression of neurotoxic astrocyte genes. As expected, *icv* co-injected M3 significantly abrogated the brain mRNA expression of all five *icv* eCIRP-induced neurotoxicity-associated astrocyte genes (C3 by 53.3%, H2-T23 by 41.8%, H2-D1 by 66.2%, GBP2 by 51.1% and PSMB8 by 59.0%) (**Figure 6C**).

**Figure 6.**
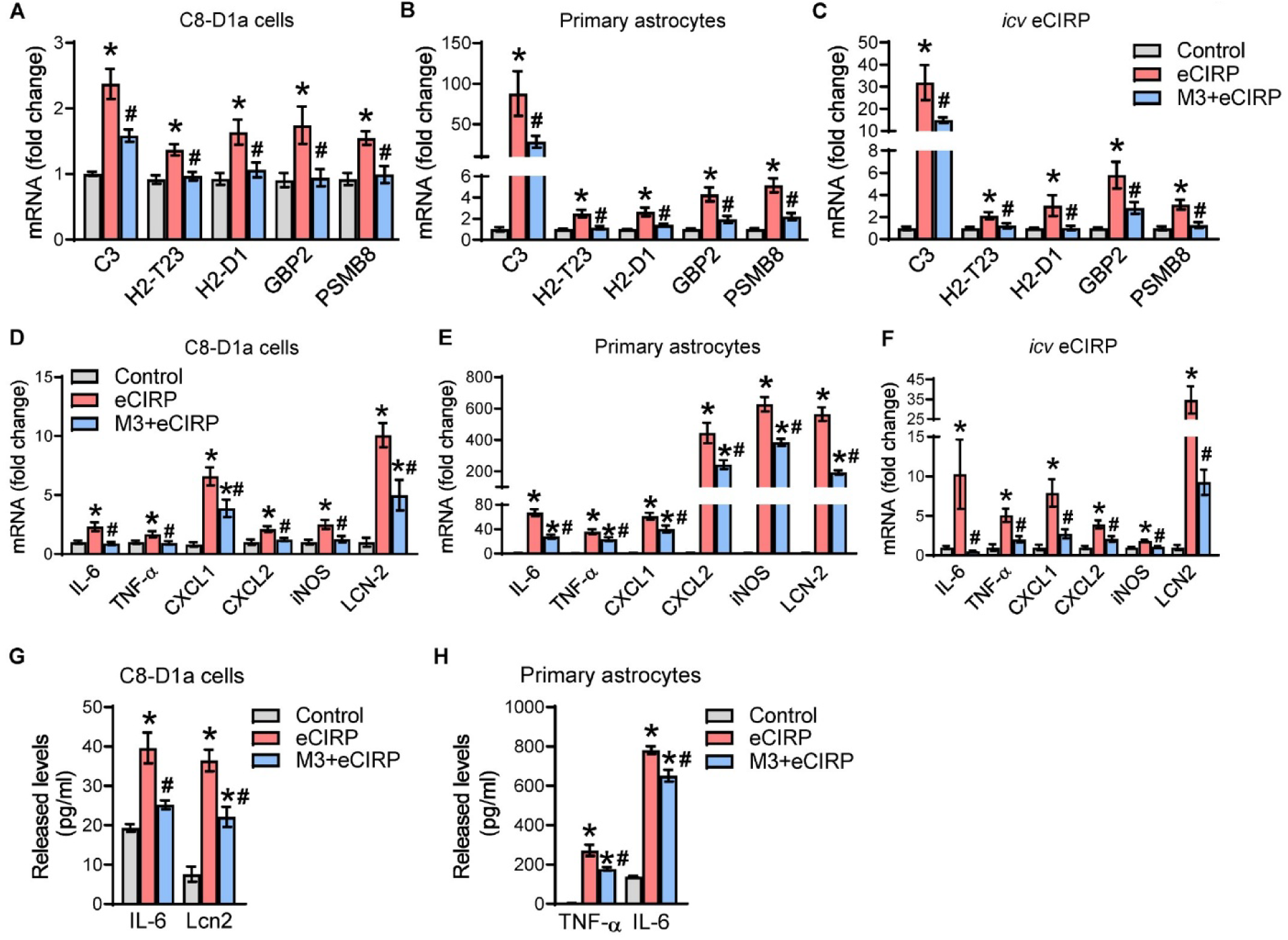
M3 attenuates eCIRP’s induction of neurotoxic astrocytes *in vitro* and *in vivo.* **(A-B)** C8-D1a cells (0.3 × 10^6^ /ml) or primary astrocytes (0.3 × 10^6^ /ml) were pretreated with 25 μg/ml M3 for 30 minutes followed by stimulation with 2.5 μg/ml eCIRP for 24 hours. RT-qPCR quantification of the relative mRNA expression of neurotoxic astrocyte signature genes C3, H2-T23, H2-D1, GBP2, and PSMB8 *in vitro* in (**A**) C8-D1a cells and (**B**) primary astrocytes. Data are expressed as means ± SEM and were compared using one-way ANOVA, Tukey’s multiple comparisons test. n = 6-8 per group; * p < 0.05 vs. control (no eCIRP), ^#^p < 0.05 vs. eCIRP alone. (**C**) 10-week-old C57BL/6 mice were *icv* co-injected with 10 mg/ml M3 or vehicle control with 1 µg eCIRP or PBS and the brains harvested 24 hours later. RT-qPCR quantification of the relative mRNA expression of the same five neurotoxic astrocyte signature genes *in vivo* in the whole brains. Data are expressed as means ± SEM and were compared using one-way ANOVA, Tukey’s multiple comparisons test. n = 8 per group; * p < 0.05 vs. control (vehicle), ^#^p < 0.05 vs. eCIRP alone. (**D-E**) C8-D1a cells (0.3 × 10^6^ /ml) or primary astrocytes (0.3 × 10^6^ /ml) were pretreated with 25 μg/ml M3 followed by stimulation with 2.5 μg/ml for 24 hours. RT-qPCR quantification of the relative mRNA expression of inflammatory cytokines (TNF-α, IL-6), chemokines (CXCL1, CXCL2), inducible nitric oxide (iNOS) and neurotoxic factor Lcn2 *in vitro* in (**D**) C8-D1a cells and (**E**) primary astrocytes. Data are expressed as means ± SEM and were compared using one-way ANOVA, Tukey’s multiple comparisons test. n = 6-8 per group; *p < 0.05 vs. control (no eCIRP), ^#^p < 0.05 vs. eCIRP alone. (**F**) C57BL/6 mice were *icv* co-injected with 10 mg/ml M3 or vehicle control with 1 µg eCIRP or PBS and the brains harvested 24 hours later. RT-qPCR quantification of the relative mRNA expression of the TNF-α, IL-6, CXCL1, CXCL2, iNOS, and Lcn2 *in vivo* in the whole brains. Data are expressed as means ± SEM and were compared using one-way ANOVA, Tukey’s multiple comparisons test. n = 8 per group; * p < 0.05 vs. control (vehicle), ^#^p < 0.05 vs. eCIRP alone. (**G-H**) C8-D1a cells (0.3 × 10 /ml) or primary astrocytes (0.3 × 10^6^ /ml) were pretreated with 25 g/ml M3 for 30 min followed by stimulation with 2.5 µg/ml eCIRP for 24 hours. Conditioned media were harvested and subjected to IL-6, TNF-α, and Lcn2 ELISA. (**G**) IL-6, and Lcn2 released in the conditioned medium of C8-D1a cells. (**H**) IL-6, and TNF-α released in the conditioned medium of primary astrocytes. Data are expressed as means ± SEM and were compared using one-way ANOVA, Tukey’s multiple comparisons test. n = 6-8 per group; *p < 0.0001 vs. no eCIRP control, ^#^p < 0.0001 vs. eCIRP alone.

M3 also effectively attenuated eCIRP-induced IL-6, TNF-α, CXCL2, and iNOS by bringing the mRNA expression levels down to control baseline in C8-D1a cells, while significantly downregulating CXCL1 by 41.3% and Lcn2 by 50.5% (**Figure 6D**). Similarly, M3 also reduced the eCIRP-induced mRNA expression of same inflammatory genes (IL-6 by 41.2%, TNF-α by 34.4%, CXCL1 by 33.9%, CXCL2 by 45.2%, iNOS by 38.7%, and Lcn2 by 65.8%) in primary astrocytes as well (**Figure 6E**). Likewise, M3 *icv*-injected in mice brains *in vivo* effectively attenuated the eCIRP-induced mRNA expression of IL-6 and iNOS down to control baseline while significantly decreasing Lcn2 by 73.4%, CXCL1 by 65.3%, TNF-α by 58.8%, and CXCL2 by 46.4% (**Figure 6F**).

Finally, M3 also significantly abrogated eCIRP-induced released IL-6 by 36.4%, and Lcn2 by 39.3% in the conditioned medium of C8-D1a cells (**Figure 6G**). Primary astrocytes pretreated with M3 also showed attenuation of eCIRP-induced released inflammatory mediators in the conditioned medium (TNF-α by 34.6% and IL-6 by 16.7%) (**Figure 6H**). Of note, the eCIRP-induced IL-6 release levels were 19.7-fold higher (**Figure 6G-H**) and Lcn2 release levels were 47.9-fold higher (**Supplementary Figure 7**) in primary microglia compared to C8-D1a cells which could explain better efficiency of M3 in C8D1a-cells in attenuating these inflammatory markers. These results show that M3 is successful in inhibiting eCIRP induction of proinflammatory and neurotoxic astrocytes *in vitro* as well as *in vivo*.

## Discussion

Elevated plasma eCIRP levels have been of potential prognostic value in a variety of inflammatory diseases such as pancreatitis, pneumonia, coronary syndrome, COVID-19, systemic sclerosis-associated interstitial lung disease (60–65), and more recently in ischemic brain injury (66). The BBB is disrupted in AD, allowing for the detection of CSF-originated AD biomarkers such as Aβ^42^, p-tau181, GFAP in the blood which are associated with cognitive decline in AD patients (50). However, the CSF and plasma eCIRP levels in AD patients have not yet been reported. eCIRP is a proinflammatory mediator and polarizes a variety of cell types towards type 1 immune responses (21, 35–40), thus mediating tissue injury in CNS and systemic pathological processes (21, 28, 67–73). While neurotoxic astrocytes appear to be pathogenic in AD/AD-tauopathy (11–14), eCIRP’s role in astrocytic neuroinflammation in AD and the involved mechanisms have yet to be investigated. Despite advances in AD drug discovery and development (74), a high unmet need remains for novel targets and effective therapeutic strategies to prevent, slow, or cure AD. This is due to modest efficacy and serious side effects associated with the recent Aβ-targeting FDA-approved drugs (75, 76). Discovering the molecular mechanisms of eCIRP associated with neurotoxic astrocytes could potentially lead to new targeted therapies for cognitive deficits in AD.

In the present study, we report for the first time that eCIRP markedly increased in the CSF and plasma of AD patients, which may have diagnostic and prognostic value in AD. eCIRP also increased in hTau.P301S mice plasma prior to as well as with AD tauopathy. We have also shown that eCIRP is associated with astrocyte reactivity in AD patients. Thus, we cogitated that released eCIRP levels from activated microglia which increase due to neuroinflammatory stress can induce neurotoxic astrocytes in AD. Indeed, we show for the first time that eCIRP is sufficient to upregulate expression of neurotoxic signature genes and inflammatory markers in astrocytes along with inducing release of proinflammatory and neurotoxic factors in vitro as well as in vivo. Herein, we report that neuroinflammatory stress releases eCIRP in brain, which binds to and activates astrocytic TREM-1 pathway to induce neurotoxic astrocytes producing C3 and inhibitory peptide M3 targeting this interaction attenuates neurotoxic astrocytes and could be a potential novel strategy to attenuate cognitive deficits in AD (**Figure 7**).

**Figure 7.**
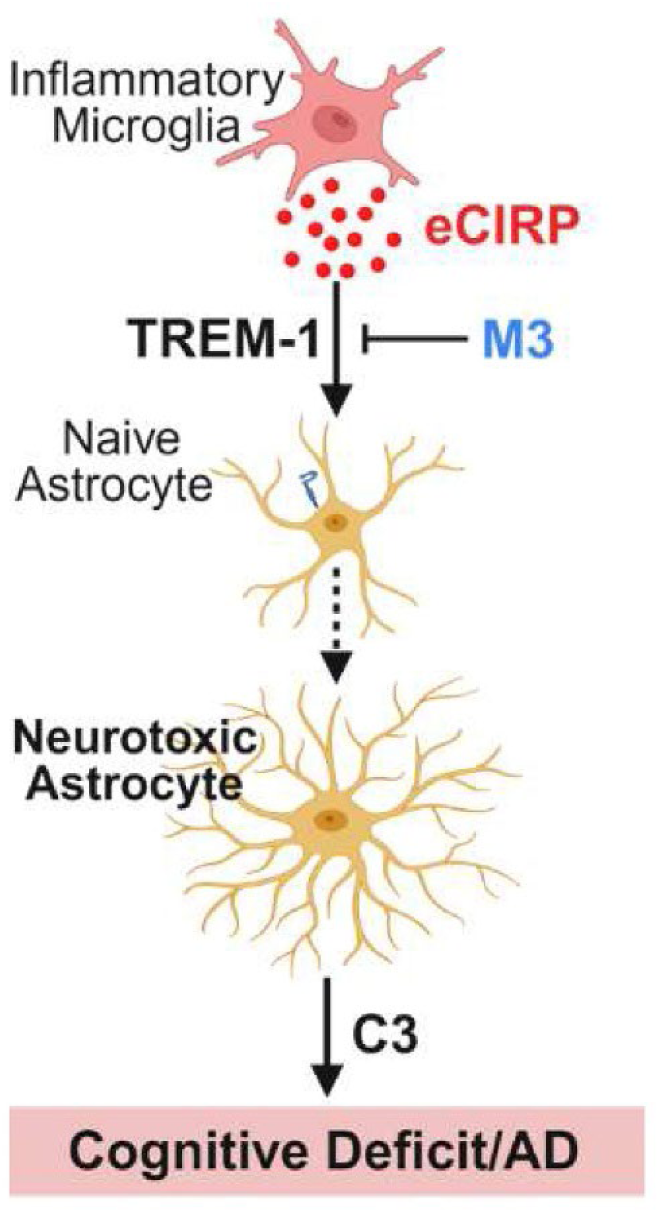
Schematic summary for eCIRP induction of neurotoxic astrocytes. Chronic neuroinflammatory stress activates microglia to release eCIRP in brain, which binds to and upregulates TREM-1 expression and activation in naïve astrocytes leading to induction of neurotoxic genes and release of proinflammatory mediators in particular C3 from neurotoxic astrocytes causing cognitive deficits in AD. The oligopeptide M3 attenuates neurotoxic astrocytes by targeting astrocytic TREM-1 activation by eCIRP and is predicted to attenuate the cognitive deficits in AD.

The eCIRP levels in plasma as well as CSF from AD patients were elevated, but the plasma eCIRP levels showed a greater increase from non-AD controls in comparison to the CSF eCIRP levels. It is important to clarify that the de-identified CSF and plasma samples tested were not from the same cohort of AD patients. In contrast to healthy aged brain AD brain is more hyperinflammatory (77), which can cause elevation of eCIRP levels in the CSF of AD patients. Of note the plasma eCIRP levels in the age-matched non-AD controls were much lower than the CSF eCIRP levels. These differences in plasma and CSF eCIRP levels could be partly explained by total blood volume being much larger than the CSF volume, causing massive dilution of eCIRP that do cross over BBB in healthy non-AD controls. Additionally, active proteases and soluble receptors present in the plasma, along with peripheral hepatic/renal clearance mechanisms could rapidly remove circulating eCIRP (25, 78). The BBB is more disrupted in AD (50) which explains more eCIRP getting in plasma from CSF despite dilution in plasma.

We also showed positive correlation between plasma eCIRP and the astrogliosis marker GFAP in a smaller cohort of AD patients. Increased plasma levels of GFAP have been strongly linked with cognitive impairment in AD patients (53, 54). Besides, the neurotoxic astrocyte marker C3 is elevated in CSF of AD patients (79), which further supports the eCIRP’s association with neurotoxic astrocytes in AD. Notably, hTau.P301S mice showed significantly lower plasma levels of eCIRP compared to AD patients. These differences between the plasma eCIRP levels in this pure tauopathy AD mouse model and the human patients could be driven by the lack of Aβ pathology in hTau.P301S mice, combined with the relative lack of widespread BBB disruption in hTau.P301S mice (80) in contrast to AD patients.

Interestingly, overexpression of intracellular CIRP (iCIRP) in human glioma cell line SHG-44, derived from a grade IV astrocytoma, was shown to have neuroprotective effects with improved cell viability under oxidation, chemical hypoxia, glucose deprivation and glutamate neurotoxicity, and upregulated EGF expression (81). In contrast, a recent study overexpressed intracellular CIRP in three human glioma cell lines U87, U251 and H4 and showed that on their coculture with human neuroblastoma SH-SY5Y cells, the neuronal beta-secretase, Aβ1-42 production and AD-like tau phosphorylation (Ser396) was increased (82). The effect of CIRP overexpression in this study was partly mediated by the downregulation of astrocytic urokinase plasminogen activator (uPA), but the molecular mechanisms remain unknown (82). Although this supports our concept that astrocytic CIRP drives AD-like neuropathy, there are clear and striking differences in functions of iCIRP and eCIRP (83, 84), which need clarification in the context of astrocytes due to contrasting data reported in above studies. Additionally, the use of oncogenic glioblastoma/astrocytoma cell lines in these studies is a serious limitation as they are significantly different in critical astrocytic features and signaling from normal astrocytic cell lines as well as from primary astrocytes.

It is well known that primary astrocytes express significantly higher levels of mature astrocytic functional markers and generally exhibit higher sensitivity and more robust physiological responses to external stimulation compared to immortalized or transformed astrocytic cell lines like C8-D1a. (85). We assessed eCIRP’s effect on both due to higher experimental reproducibility and stability associated with C8-D1a cells compared to batch-to-batch donor variability and potential microglial/oligodendrocyte contamination risk associated with primary astrocytes which could offset their high physiological relevance and sensitivity benefits. Accordingly, primary astrocytes showed more sensitivity to eCIRP stimulation compared to C8-D1a cells in our study with more robust neurotoxic signature and higher levels of expression and release of inflammatory and neurotoxic mediators such as IL-6 and Lcn2.

In particular, eCIRP increased cellular as well as released levels of astrocytic C3, a neurotoxic marker involved in neuroinflammation-associated neurodegeneration (79). Targeting TREM-1 genetically (TREM-1^−/−^ astrocytes) or pharmacologically (TREM-1 inhibitory peptide LP17 and eCIRP/TREM-1 specific inhibitory peptide M3) attenuated eCIRP’s induction of neurotoxic astrocytes. eCIRP interaction with TREM-1 and its inhibition by the peptide M3 has been well established by our group in a variety of disease settings (40, 45–49, 86–90). Neurotoxic astrocytes depend on NFκB (11) and TREM-1 signaling induces downstream NFκB (91, 92). We showed that eCIRP activated astrocytic TREM-1 expression and downstream signaling to NFκB for neurotoxic astrocyte induction. eCIRP directly induced TREM-1 downstream Syk phosphorylation and NFκB induction in C8-D1a cells. Another recent study showed that the transcription factor Yin Yang-1 binds to the TREM-1 promoter to activate NF-κB signaling in LPS-treated mouse astrocytes in the context of chronic stress-induced depression, which supports our data (93). Therefore, eCIRP induction of NFκB in primary astrocytes fits upstream of neurotoxic astrocyte generation.

Neurotoxic astrocytes are also critically regulated by the nuclear factor erythroid 2-related factor 2 (Nrf2) pathways (14). We will need to further identify the signaling molecules downstream from TREM-1 leading to NFκB activation, which in turn suppresses Nrf2. In future, we will dissect whether Nfr2 is involved in eCIRP’s induction of neurotoxic astrocytes and verify the critical roles of NFκB and Nfr2 pathways in AD pathogenesis *in vivo*. We expect eCIRP-induced NFκB activation to downregulate Nrf2. Other TREM-1 pathways, such as PI3K/AKT and JAK/STAT pathways (42) may also play role in eCIRP induction of neurotoxic astrocytes via TREM-1/NFκB axis and would need further studies to access their involvement.

Current study showed that eCIRP activates TREM-1 signaling and upregulates NFκB, leading to the expression of neurotoxic signature, in particular C3. Astrocytic NFκB activation causes release of C3 which in turn interacts with neuronal and microglial C3a receptor (C3aR) to respectively alter cognitive function and promote synaptic pruning via microglial phagocytosis (94–96). C3 plays critical role in neuroinflammation and neurodegeneration via C3aR (79, 97). Future studies to evaluate C3 as a critical effector by which eCIRP-induced neurotoxic astrocytes cause neuronal and microglial dysfunction are warranted.

Of note most studies for neurotoxic astrocytes in AD have been focused on amyloid-based in vivo AD models (12, 98, 99). In light of the earlier data from our lab showing eCIRP association with AD-tauopathy, it would be particularly adequate to study eCIRP-induced neuroinflammation in the context of tau pathology (100–102). We also demonstrated that M3 attenuates eCIRP-induced neurotoxic astrocytes by specifically targeting eCIRP/TREM-1 interaction. As targeting reactive astrocytes as well as blocking C3 improves cognition and behavioral deficits (79), it would be imperative to investigate if M3 can improve the cognition deficits in AD mouse models of tauopathy such as Tau P301S or 3xTg mice with combined amyloid and tauopathy.

In summary, we show that eCIRP is elevated in CSF and plasma in AD patients and strongly correlates with astrocyte reactivity, highlighting the clinical relevance of our findings. eCIRP directly induced neurotoxic astrocytes via TREM-1, and M3 – a CIRP-derived 7-mer peptide – effectively attenuated eCIRP-induced neurotoxic astrocytes by blocking eCIRP’s activation of the TREM-1 pathway. Taken together, the current study strongly indicates that eCIRP induces neurotoxic astrocytes via TREM-1 and that M3 reduces neurotoxic astrocytes and is thus expected to attenuate AD-associated cognitive deficits.

## Methods

### Sex as a biological variable

Considering gender differences in Alzheimer’s disease pathology, human plasma samples from both males and females (38% males 62% females reflecting the sex differences in AD) were included in the study to allow exploratory comparisons using sex as an independent variable. The human findings are expected to be relevant for both males and females. Our *in vitro* data used the whole litter to generate primary astrocytes, thus including both male and female pups. To exclude the effects of sex-specific AD differences on in vivo mechanistic studies, only male C57BL/6 (for icv injections) and hTau.P301S (for plasma) mice were used in in this study to generate reliable and consistent findings. It is unknown whether the findings are relevant to female mice.

### Human plasma and CSF samples

Plasma samples were purchased from the deidentified age-and sex-matched unaffected human subjects and AD patients from Precision Biospecimen Solutions (Bethesda, MD). There are 14 males (37.84%) and 23 females (62.16%) in each group with the age range 63-88 years. There was no significant difference in median ages of control group (77 years) vs. AD group (79 years). Control samples were non-reactive in viral testing for HIV, HBsAg and HCV. The demographic data including clinical CDR and MMSE test scores, cognitive change status and diagnostic imaging results were provided for the AD patients. The de-identified CSF samples from unaffected healthy elderly controls and AD patients were obtained from the biorepository at Litwin-Zucker Center for Alzheimer’s Disease Research and diagnosed according to current criteria, which includes extensive annual cognitive testing in addition to structural imaging and ApoE genotyping (52, 103, 104). The shared specimens originate from a well-characterized longitudinal cohort of subjects from Alzheimer’s Disease Neuroimaging Initiative (62).

### CIRP and GFAP ELISA

Human CIRP ELISA kit (catalog no. CSB-EL005440HU, American Research Products) and human GFAP ELISA kit (catalog no. NBP3-11815, Novus Biologicals) were used for quantifying CSF eCIRP, plasma eCIRP and plasma GFAP concentrations as per manufacturer’s protocol. Mouse CIRP ELISA kit (catalog no. EM6504, FineTest) was used to measure eCIRP in plasma samples from hTau.P301S mice.

### Recombinant proteins and peptides

Recombinant mouse CIRP (referred to as eCIRP) was prepared in-house, with quality control measures performed as previously described (21). M3 (RGFFRGG) and LP17 (LQVTDSGLYRCVIYHPP) peptides were synthesized by GenScript USA Inc., purified to > 95% by HPLC, and provided as a lyophilized powder. The peptide was resuspended in sterile PBS at desired concentration prior to treatment of cells or mice.

### C8-D1a cell culture and treatment

Mouse astrocyte type I clone (C8-D1a) cells were purchased from ATCC (catalog no. CRL-2541) and were cultured in DMEM supplemented with 10% FBS and 100 U/mL penicillin/streptomycin in a humidified atmosphere at 37° C, 5% CO2. C8-D1a cells were detached with 0.25% trypsin-EDTA, passaged 2-3 times weekly, and seeded at a density of 0.3 × 10^6^ cells per well for experiments. After resting overnight, media was exchanged to fresh DMEM and treated with PBS or eCIRP (doses specified in each experiment) for 24 hours (or duration specified in each experiment) or pretreated with 30 minutes with LP17 (80 μg/ml) or M3 (25 μg/ml) for 30 min followed by stimulation with 2.5 µg/ml eCIRP for 24 hours. Conditioned media were harvested and cells were lysed for protein or mRNA extraction.

### Experimental animals

Male 8-week-old C57BL/6 mice were purchased from Charles River Laboratories. Mice were allowed to acclimate for 1 week before use in experiments. The TREM-1^−/−^ mice (Trem1^tm1(KOMP)Vlcg^) generated by the trans-NIH Knockout Mouse Project (KOMP) were obtained from the KOMP Repository, University of California and maintained. Mice were housed at the Center for Comparative Physiology at the Feinstein Institutes for Medical Research in a temperature-controlled room under 12-hour light/12-hour dark cycles and given standard laboratory food and water ad libitum. For primary microglia isolation, breeding triads were closely monitored to precisely record the pups date of birth. House-bred C57BL/6 and TREM-1^−/−^neonatal mouse pups aged 0-3 days (P0-P3) were utilized for primary astrocyte isolation. All experiments were performed following the NIH guidelines for experimental animals.

### Isolation and purification, culture and treatment of primary murine astrocytes

Primary astrocytes were isolated from C57BL/6 or TREM-1^−/−^ P0-P3 neonatal mouse pups as previously described (105). Astrocytes were magnetically purified by positive selection using the anti-ACSA-2 magnetic microbeads (catalog no. 130-097-679, Miltenyi Biotech). Briefly, whole brains were harvested in Hank’s balanced salt solution and dissociated with Neural Tissue Dissociation Kit (P) (catalog no. 130-092-628, Miltenyi Biotec) per the manufacturer’s protocol. The purity of the astrocytes was assessed by staining of the cells with APC-GLAST antibody (catalog no.130-123-641, Miltenyi Biotec) using a BD LSRFortessa flow cytometer (BD Biosciences). The astrocytes were cultured in Gibco Neurobasal™ Medium (catalog no. 21103-049, Thermo Fisher Scientific) supplemented with 1X serum free B-27 (catalog no. 17504-044, Thermo Fisher Scientific) and 5 ng/ml HB-EGF (catalog no. 100-47, Peprotech) and seeded in T-175 culture flasks. Thereafter, culture medium was changed twice weekly. Astrocytes were recovered from flasks at DIV 10-21 by detacheding with 0.25% trypsin-EDTA, centrifuged at 400g for 10 min, and resuspended in complete DMEM prior to seeding at a density of 0.3 × 10^6^ cells per well for experiments in tissue culture plates and rested overnight for at least 16 h before treatment. Primary astrocytes were treated with PBS or eCIRP (doses specified in each experiment) for 24 hours (or duration specified in each experiment) or pretreated with 30 minutes with LP17 (80 μg/ml) or M3 (25 μg/ml) for 30 min followed by stimulation with 2.5 µg/ml eCIRP for 24 hours. Conditioned media were harvested and cells were lysed for protein or mRNA extraction.

### Intracerebroventricular injection

Male C57BL/6 mice were subjected to intracerebro-ventricular (icv) injection as previously described (106). Briefly, mice were anesthesised with isoflurane, scalp shaved, disinfected, placed in the stereotactic apparatus and a small burr hole was drilled in the skull with stereotaxic Dremel using Bregma coordinates at AP 0.34 mm, lateral 1.0 mm and vertical 2.2 mm. 2 µL eCIRP (1 µg) or PBS was injected using a 33-gauge Hamilton syringe attached to the stereotaxic injector (Stoelting) at a rate of 0.2 µL per minute. The needle was left in place for 5 minutes, then was slowly withdrawn and skin was closed with silk suture. Mice were sacrificed 24 hours post-icv, brain tissues were collected and frozen in liquid nitrogen before storage at −80° C.

### Real-time qPCR

Total RNA was isolated from cells or whole brains with RNAspin Mini kit (Cytiva). An equal amount (300 ng – 1 µg) of RNA was reverse-transcribed into cDNA using murine leukemia virus reverse transcriptase enzyme (Thermo Fisher Scientific). qPCR was performed from the diluted cDNA with specific forward and reverse primers (**Supplementary Table 1**) and SYBR green PCR Master Mix (catalog no. 4312704, Applied Biosystems) using QuantStudio3 real-time thermocycler (Applied Biosystems). Mouse β-actin served as an internal control gene for normalization. The relative mRNA expression was expressed as the fold change compared to the control quantified using comparative cycle threshold (CT) method of 2^(-ΔΔCT)^.

### ELISA and Proteome profiling for released inflammatory mediators

The levels of specific inflammatory mediators released in the condition medium of C8-D1a cells or primary astrocytes were quantified using BD OptEIA™ Mouse IL-6 ELISA (catalog no. 555240, BD Biosciences), and BD OptEIA™ Mouse TNFα ELISA (catalog no. 558534, BD Biosciences), Mouse Lcn-2 ELISA (catalog no. MLCN20, R&D Systems), and Mouse C3 ELISA (catalog no. OKCD06290, Aviva) as per manufacturer’s instructions. The levels of 111 inflammatory mediators released in the condition medium of primary astrocytes were quantified using Proteome Profiler Mouse XL Cytokine Array (catalog no. ARY028, R&D Systems) as per manufacturer’s instructions.

### Western blot assays

Whole-cell proteins were extracted with ice cold RIPA lysis buffer (10 mM Tris buffered Saline (TBS) pH 7.5, containing phenylmethylsulfonyl fluoride, Na-orthovanadate, protease and phosphatase inhibitor cocktails (Thermo fisher Scientific). Protein concentration was measured via DC protein assay (catalog no. 5000111, Bio-Rad). Proteins were separated via electrophoresis using NuPAGE™ 4%–12% Bis-Tris gels (Invitrogen) and wet-transferred onto nitrocellulose membranes. Membranes were blocked in 0.1% casein in TBS for 1 hour at room temperature and then incubated in blocking solution containing 0.1% Tween-20 overnight at 4°C with primary antibodies according to the manufacturer’s recommendations. The primary antibodies used were C3 (catalog no. ab97462, Abcam, 1:1000), S100A10 (catalog no. AF2377, R&D Systems, 1:2000), TREM-1 (catalog no. ab104413, Abcam, 1:1000), phospho-Syk (Tyr525, Tyr526) (clone F.724.5, catalog no. MA5-14918, Thermo Fischer Scientific, 1:000), Syk (clone D3Z1E, catalog no. 13198, Cell Signaling, 1:000), and NFκB p65 (clone D14E12, catalog no. 8242, Cell Signaling, 1:1000). After the blots were incubated with primary antibodies, the membranes were washed 3 times, and the blots were incubated with the corresponding infrared dye-labeled secondary antibodies (anti–mouse IgG, catalog 926-68070; anti–rabbit IgG, catalog 926-32211; Li-Cor Biosciences) for detection. Membranes were imaged for bands using an Odyssey Clx Imaging system with Image Studio 5.2 software (Li-Cor Biosciences). After target band imaging, membranes were incubated to measure endogenous loading control with β-actin antibody (catalog no. A5441, Sigma-Aldrich, 1:10000) and bands imaged in the same manner. Densitometric quantification was done using NIH ImageJ.

### Confocal microscopy

Primary astrocytes or C8-D1a cells were blocked with 5% normal horse serum for 1 hour and stained with for goat anti-mouse TREM-1 (catalog no. AF1187, R&D Systems, 1:100) or rabbit anti-mouse GFAP (catalog no. ab68428, Abcam, 1:100) primary Abs diluted in 1% horse serum for 2 hours at room temperature or overnight at 4°C. The cells were washed in PBS and were incubated in the dark with diluted Cy5-conjugated AffiniPure donkey anti-goat IgG (code 705-175-147; Jackson ImmunoResearch Laboratories) or Cy3-conjugated AffiniPure F(ab′)2 donkey anti-rabbit IgG (code 711-166-152; Jackson ImmunoResearch Laboratories) secondary Abs for 1 hour. After an additional washing step, slides were mounted immediately on Vectashield mounting medium with Hoechst (Vector Laboratories). Slides were imaged on a Zeiss LSM880 scanning confocal microscope (Carl Zeiss Imaging, Germany). Z-stack images were obtained using a 63x objective. Images of 2-3 different well regions were obtained. Images were analyzed using ZenBlue software (Zeiss).

### Flow cytometry

C8-D1a cells or primary astrocytes were treated with 2.5 μg/mL eCIRP for 24 hours. After 24 hours, cells were detached for 15 min in cold PBS containing 1% FBS. Cells were then stained with APC anti-mouse TREM-1 Ab (clone 174031, catalog no. FAB1187A, R&D Systems). Unstained cells were used as a negative control to establish the flow cytometer voltage setting. Acquisition was performed on 10,000 events using a BD FACSymphony flow cytometer (BD Biosciences). FCS files were exported and analyzed in FlowJo 10.9.0 (BD Biosciences).

### Statistics

Data represented in the figures are expressed as mean ± SEM and were compared by Mann-Whitney test for 2 groups for human plasma samples or unpaired 2-tailed Student’s *t* test for 2 groups or Multiple group *t-*test or 1-way ANOVA using post-hoc Tukey’s multiple comparisons test or two-way ANOVA using Sidak’s multiple comparisons test or Mixed-effects model (REML) with Tukey’s multiple comparisons test for data with missing values. Correlation was analyzed using Linear Regression and Spearman’s Correlation. Differences in values were considered significant if *P* was less than or equal to 0.05. Data analysis was carried out using GraphPad Prism graphing and statistical software (GraphPad Software Inc., La Jolla, CA).

### Study approval

The present studies using live animals were reviewed and approved by the Institutional Animal Care and Use Committee of the Feinstein Institutes for Medical Research.

## Data availability

All data generated or analyzed during this study are included in this manuscript.

## Author Contributions

AS designed the experiments. AS and DA conducted all in vitro experiments, acquired, and analyzed the data. DL performed in vivo *icv* injections. AS interpreted data and wrote and edited the manuscript. PM provided reagents (hTau.P301S mice plasma) and critical input in some experimental designs. PW critically reviewed the manuscript. AS and PW conceived the idea. AS and PW supervised the project. All authors read and approved the final manuscript.

## Acknowledgements

The authors sincerely acknowledge that Dr. Monowar Aziz played a significant role in the discovery of TREM-1 as a receptor of eCIRP and the development of M3 as anti-eCIRP peptide. Those discoveries provide a strong foundation for the current work. The authors acknowledge Dr. Wayne Chaung for assistance with animal colony maintenance and Dr. Yongchan Lee for his technical assistance with the confocal microscopic study. We also thank former research assistant Ezgi Sari for technical assistance with proteome profiling. We are thankful to our colleague, Dr. Sangeeta Chavan, for sharing aliquots of some of the plasma samples from AD patients and control subjects for eCIRP ELISA. We acknowledge the use of BioRender in creating schematics.

