## Supplemental Materials for "Extracellular CIRP induces neurotoxic astrocytes via TREM-1 in Alzheimer’s disease"

**Supplementary Data Figures**

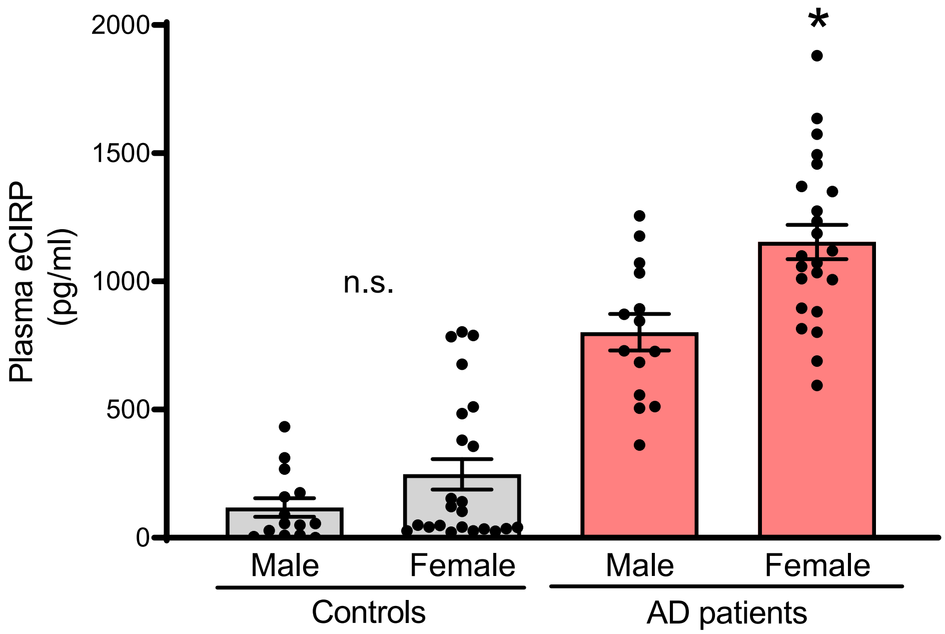


**Supplementary Figure 1. eCIRP levels are higher in female AD patients.** The plasma levels of eCIRP were measured by ELISA in AD patient and age- and sex-matched unaffected control samples obtained from Precision Biospecimen Solutions (Bethesda, MD). There are 14 males (37.84%) and 23 females (62.16%) per group with the age range 63-88 years and no significant difference in median age. Experiments were performed at least 3 times, and all data were analyzed. Data are expressed as means ± SEM and were compared using Mixed-effects model (REML) with Sidak's multiple comparisons test. n = 14 males per group and 23 females per group; n.s. $p = 0.26$ for male vs female control subjects,*$p = 0.0005$ for male vs. female AD patients.

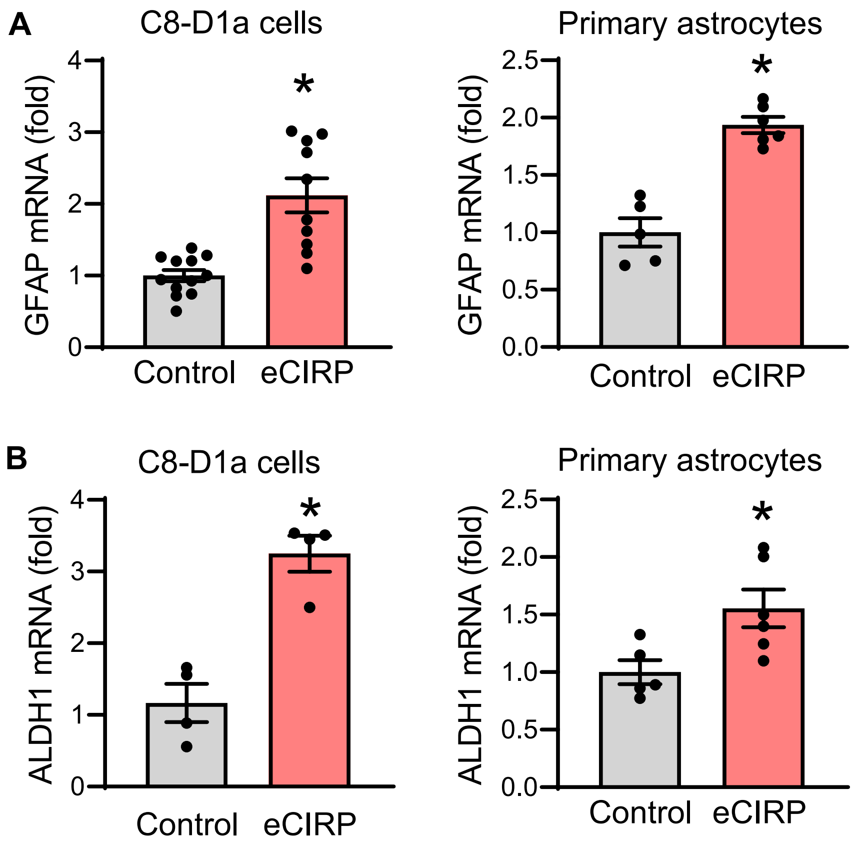


**Supplementary Figure 2. Astrocyte marker genes GFAP and ALDH1 are expressed by magnetically purified primary astrocytes which are increased by eCIRP stimulation.** C8-D1a cells ($0.3 \times 10^6$/ml) or magnetically isolated primary astrocytes ($0.3 \times 10^6$/ml) were treated with 2.5 µg/ml eCIRP for 24 hours and subjected to RNA extraction. RT-qPCR quantification of the relative mRNA expression of (**A**) GFAP and (**B**) ALDH1 in C8-D1a cells and primary astrocytes as indicated. Data are expressed as means ± SEM and were compared using Unpaired *t*-test. n = 4-10 per group and * $p < 0.05$ vs. control (no eCIRP).

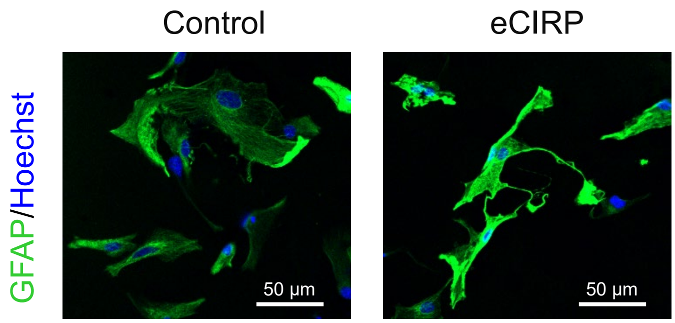


**Supplementary Figure 3. eCIRP increases GFAP protein levels in primary astrocytes.**

Magnetically isolated primary astrocytes (0.3 × $10^6$/ml) were treated with 2.5 µg/ml eCIRP for 24 hours and subjected to staining with GFAP for confocal microscopy Representative confocal microscopy images for GFAP protein expression in control and eCIRP stimulated primary astrocytes stained for Hoechst (blue), and GFAP (green). Magnification 630x, scale bar, 50 µM.

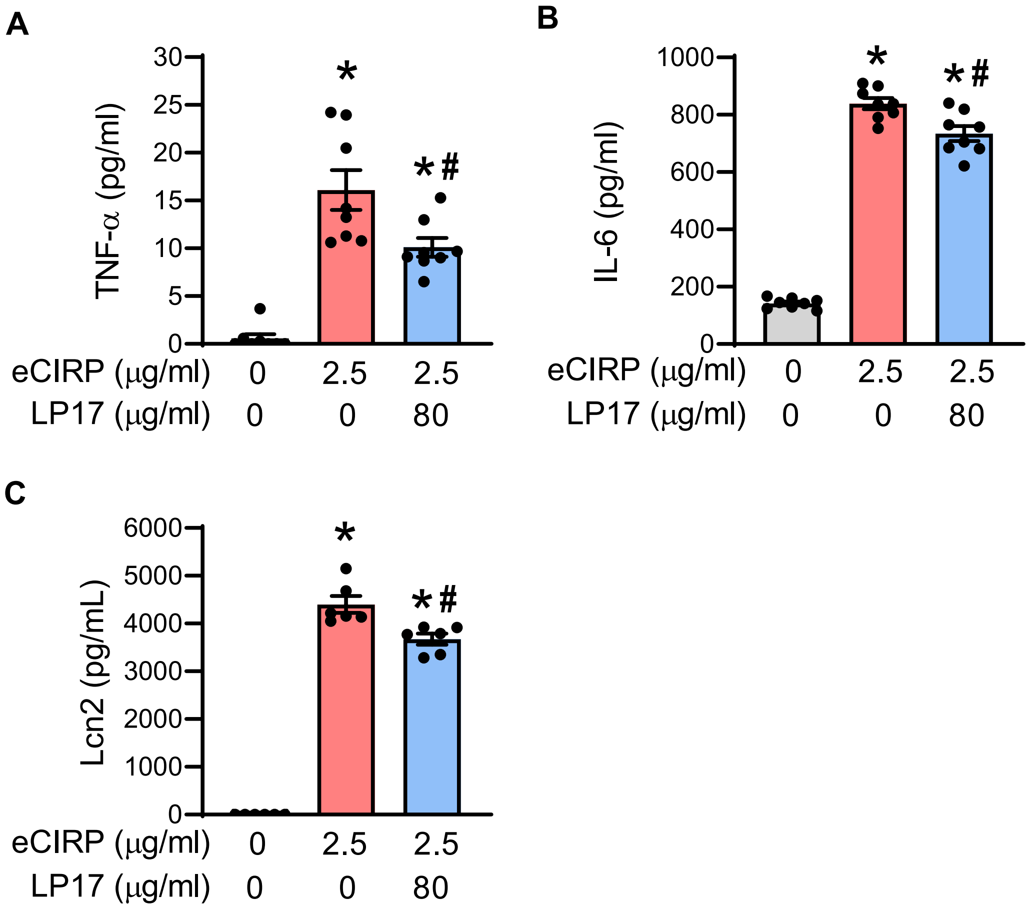


**Supplementary Figure 4. LP17 attenuates eCIRP-induced release of inflammatory mediators in primary astrocytes.** Primary astrocytes ($0.3 \times 10^6$/ml) were pretreated with 80 μg/ml LP17 (TREM-1 inhibitor peptide) for 30 min followed by stimulation with 2.5 μg/ml eCIRP for 24 hours. Conditioned media were harvested and released inflammatory mediators TNF-α, IL-6, and Lcn2 quantified by ELISA. Data are expressed as means ± SEM and were compared using one-way ANOVA with Tukey's multiple comparisons test. n = 6-8 per group; *$p < 0.05$ vs. no eCIRP control, $^{\#}p < 0.05$ vs. eCIRP alone.

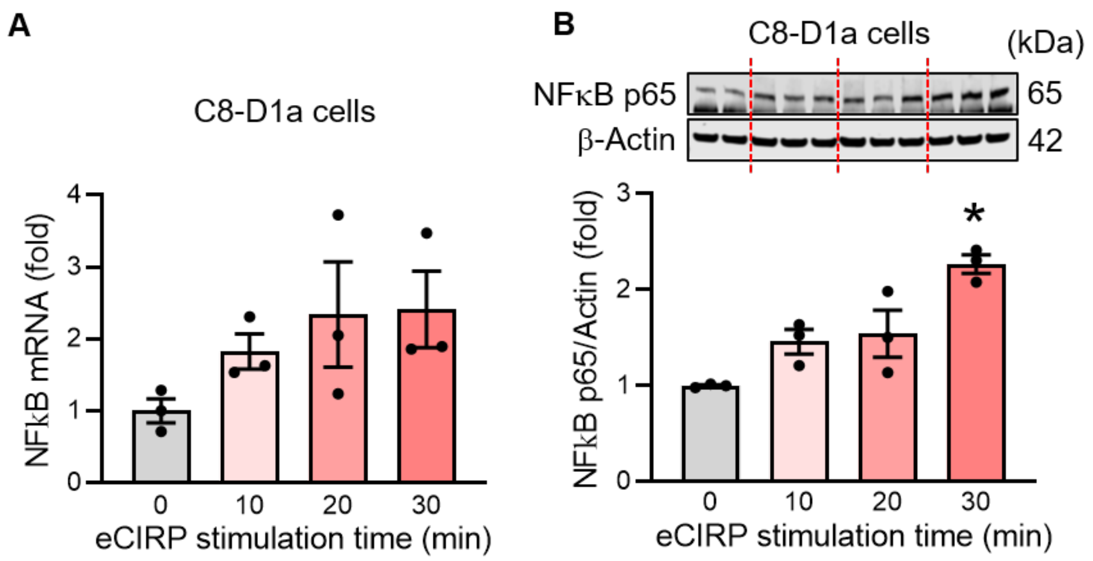


**Supplementary Figure 5. eCIRP induces early NFκB expression in C8-D1a cells in time-dependent manner.** C8-D1a cells ($0.3 \times 10^6$/ml) were treated with 2.5 µg/ml eCIRP for 0, 10, 20 and 30 minutes and subjected to RNA or protein extraction for assessing NFκB expression. (**A**) RT-qPCR quantification of the relative mRNA expression of NFκB in C8-D1a cells at indicated times. Data are expressed as means ± SEM and were compared using one-way ANOVA, Tukey's multiple comparisons test. n = 3 per group. Data not significant. (**B**) Representative Western blot images and bar graph from the densitometric analysis of NFκB p65 normalized to β-actin protein. The dotted lines on the Western blot images reflect different samples from the corresponding groups shown in the bar graph below. Data are expressed as means ± SEM and were compared using (**D**) one-way ANOVA, Tukey's multiple comparisons test. n = 3 per group; *$p < 0.05$ vs. no eCIRP control.

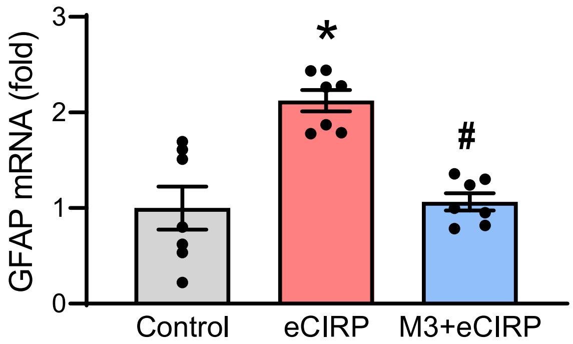


**Supplementary Figure 6. M3 attenuates eCIRP-induced GFAP mRNA expression in primary astrocytes.** Primary astrocytes ($0.3 \times 10^6$/ml) were pretreated with 25 µg/ml M3 peptide for 30 minutes followed by stimulation with 2.5 µg/ml eCIRP for 24 hours and subjected to RNA extraction. RT-qPCR quantification of the relative mRNA expression of GFAP in primary astrocytes. Data are expressed as means ± SEM and were compared using one-way ANOVA, Tukey's multiple comparisons test. n = 7 per group; * $p < 0.05$ vs. control (no eCIRP) and $^{\#}$ $p < 0.05$ vs. eCIRP alone.

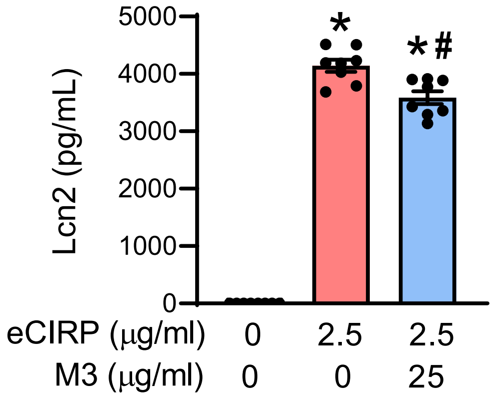


**Supplementary Figure 7. M3 attenuates eCIRP-induced Lcn2 in primary astrocytes.**

Primary astrocytes ($0.3 \times 10^6$/ml) were pretreated with 25 μg/ml M3 peptide for 30 min followed by stimulation with 2.5 μg/ml eCIRP for 24 hours. Lcn2 levels released in the conditioned media were quantified by ELISA. Data are expressed as means ± SEM and were compared using one-way ANOVA with Tukey's multiple comparisons test. n = 6-8 per group; *$p < 0.05$ vs. no eCIRP control, $^{\#}p < 0.05$ vs. eCIRP alone.

Supplementary Table 1. Primer sequences used for RT-qPCR in this study

| Target | Forward primer (5’-3’) | Reverse primer (5’-3’) |
|---|---|---|
| C3 | CCAGCTCCCCATTAGCTCTG | GCACTTGCCTCTTTAGGAAGTC |
| H2-T23 | GGACCGCGAATGACATAGC | GCACCTCAGGGTGACTTCAT |
| H2-D1 | TCCGAGATTGTAAAGCGTGAAGA | ACAGGGCAGTGCAGGGATAG |
| GBP2 | GGGGTCACTGTCTGACCACT | GGGAAACCTGGGATGAGATT |
| PSMB8 | CAGTCCTGAAGAGGCCTACG | CACTTTCACCCAACCGTCTT |
| S100A10 | CCTCTGGCTGTGGACAAAAT | CTGCTCACAAGAAGCAGTGG |
| CLCF1 | CTTCAATCCTCCTCGACTGG | TACGTCGGAGTTCAGCTGTG |
| EMP1 | GAGACACTGGCCAGAAAAGC | TAAAAGGCAAGGGAATGCAC |
| SLC10A6 | GCTTCGGTGGTATGATGCTT | CCACAGGCTTTTCTGGTGAT |
| B3GNT5 | CGTGGGGCAATGAGAACTAT | CCCAGCTGAACTGAAGAAGG |
| GFAP | GCCACCAGTAACATGCAAGA | GGCGATAGTCGTTAGCTTCG |
| NFκB2 | TGCTGATGGCACAGGACGAGAA | GTTGATGACGCCGAGGTACTGA |
| TREM-1 | ACCGCAGTGGGCTTGGGTAGGG | GAGGAAGGCTGGGCTCTGGGGACT |
| IL-6 | CCGGAGAGGAGACTTCACAG | CAGAATTGCCATTGCACAAC |
| TNF-α | AGACCCTCACACTCAGATCATCTTC | TTG CTACGACGTGGGCTACA |
| CXCL-1 | GCTGGGATTCACCTCAAGAA | ACAGGTGCCATCAGAGCAGT |
| CXCL-2 | CCCTGGTTCAGAAAATCATCCA | GCTCCTCCTTTCCAGGTCAGT |
| iNOS | GCAGGTCGAGGACTATTTCTTTCA | GAGCACGCTGAGTACCTCATTG |
| Lcn2 | CTCAGAACTTGATCCCTGCC | TCCTTGAGGCCCAGAGACTT |
| Aldh1l1 | AGCAGAGGCCATTCACAACT | GCCACCAGTCCTGAAGTGTT |
| β-actin | CGTGAAAAGATGACCCAGATCA | TGGTACGACCAGAGGCATACAG |
